# Drop-In Biofuel production by using fatty acid photodecarboxylase from *Chlorella variabilis* in the oleaginous yeast *Yarrowia lipolytica*

**DOI:** 10.1101/468876

**Authors:** Stefan Bruder, Eva Johanna Moldenhauer, Robert Denis Lemke, Rodrigo Ledesma-Amaro, Johannes Kabisch

**Affiliations:** Computer-aided Synthetic Biology, Technische Universität Darmstadt, Darmstadt, Germany; Department of Bioengineering and Imperial College Centre for Synthetic Biology, Imperial College London, London, United Kingdom

**Author notes:** **Correspondence:** Johannes Kabisch, Computer-aided Synthetic Biology, Technische Universität Darmstadt, Schnittspahnstr. 12, 64287 Darmstadt, Germany. **E-mail:**.

**Keywords:** Drop-in biofuels, clean fuels, microbial biodiesel, fatty acid photodecarboxylase, hydrocarbons, alkane, alkene, oleaginous yeast, *Yarrowia lipolytica*

## Abstract

**Background:** Oleaginous yeasts are potent hosts for the renewable production of lipids and harbor great potential for derived products, such as biofuels. Several promising processes have been described that produce hydrocarbon drop-in biofuels based on fatty acid decarboxylation and fatty aldehyde decarbonylation. Unfortunately, besides fatty aldehyde toxicity and high reactivity, the most investigated enzyme, aldehyde-deformylating oxygenase, shows unfavorable catalytic properties which hindered high yields in previous metabolic engineering approaches.

**Results:** To demonstrate an alternative alkane production pathway for oleaginous yeasts, we describe the production of diesel-like, odd-chain alkanes and alkenes, by heterologously expressing a recently discovered light-driven oxidase from *Chlorella variabilis* (CvFAP) in *Yarrowia lipolytica.* Initial experiments showed that only strains engineered to have an increased pool of free fatty acids showed to be susceptible to sufficient decarboxylation. Providing these strains with glucose and light in a synthetic medium resulted in titers of 10.9 mg/L of hydrocarbons. Using custom 3D printed labware for lighting bioreactors, and an automated pulsed glycerol fed-batch strategy, intracellular titers of 58.7 mg/L were achieved.

**Conclusions:** Oleaginous yeasts such as *Yarrowia lipolytica* can transform renewable resources such as glycerol into fatty acids and lipids. By heterologously expressing a fatty acid photodecarboxylase from the algae *Chlorella variabilis* hydrocarbons were produced in several scales from microwell plate to 400 ml bioreactors. The developed bioprocess shows a route to the renewable production of hydrocarbons for a variety of applications ranging from representing a substrate for further enzymatic or chemical modification or as a drop-in biofuel blend.

**Short abstract:** Oleaginous yeasts are potent hosts for the renewable production of lipids, fatty acids and derived products such as biofuels. Here, we describe, the production of odd-numbered alkanes and alkenes with a length of 17 and 15 carbons by expression of a fatty acid photodecarboxylase (CvFAP) from *Chlorella variabilis* in different *Yarrowia lipolytica* strains under different regimes of blue light exposure in several scales from microwell plate to 400 ml bioreactors.

## 1. Background

Modern human society is built upon readily available hydrocarbons, currently predominantly derived from fossil resources. The depletion of these as well as the adverse effects of their intense utilization have led to a variety of global challenges [1]. A concept to counteract these, is to shift towards biobased processes by finding novel and drop-in alternatives produced on the basis of renewable resources. One such alternative are so called drop-in biofuels which are substantially similar to current fuels and are not associated with some of the drawbacks of first generation biofuels like ethanol or fatty acid methyl esters [1].

In recent years a variety of enzymes for the microbial production of hydrocarbons have been discovered and exploited. The most prominently among these is the pair formed by the acyl-ACP reductase (AAR) and the decarbonylating aldehyde-deformylating oxygenase (ADO) discovered in hydrocarbon producing cyanobacteria and expressed in *E. coli* by Schirmer *et al.*, 2010. Following this first proof of concept, the route from fatty acids to hydrocarbons was optimized and transferred to SCO (single cell oil) accumulating organisms.

Oleaginous yeasts are arbitrarily defined as being able to accumulate more than 20% of their cell dry weight (cdw) as lipids. Among these, the yeast *Yarrowia lipolytica* is well exploited in respect to genetic amenability and frequently used for industrial applications [2].

The ability to produce large amounts of lipids makes it an attractive host for fatty acid-derived biofuels. Thus, the above-described pathways for hydrocarbon formation have been adapted to *Y. lipolytica* by Xu *et al.*, 2016 [3]. Fig. 1 A summarizes different strategies for fatty acid-derived hydrocarbon formation with yeasts. A more recent publication identified a promiscuous activity of an algal photoenzyme [4]. This glucose-methanol-choline (GMC) oxidoreductase termed fatty acid photodecarboxylase (FAP) was found in both, *Chlorella variabilis* (CvFAP) and *Chlorella reinhardtii* (CrFAP).

**Figure 1.**
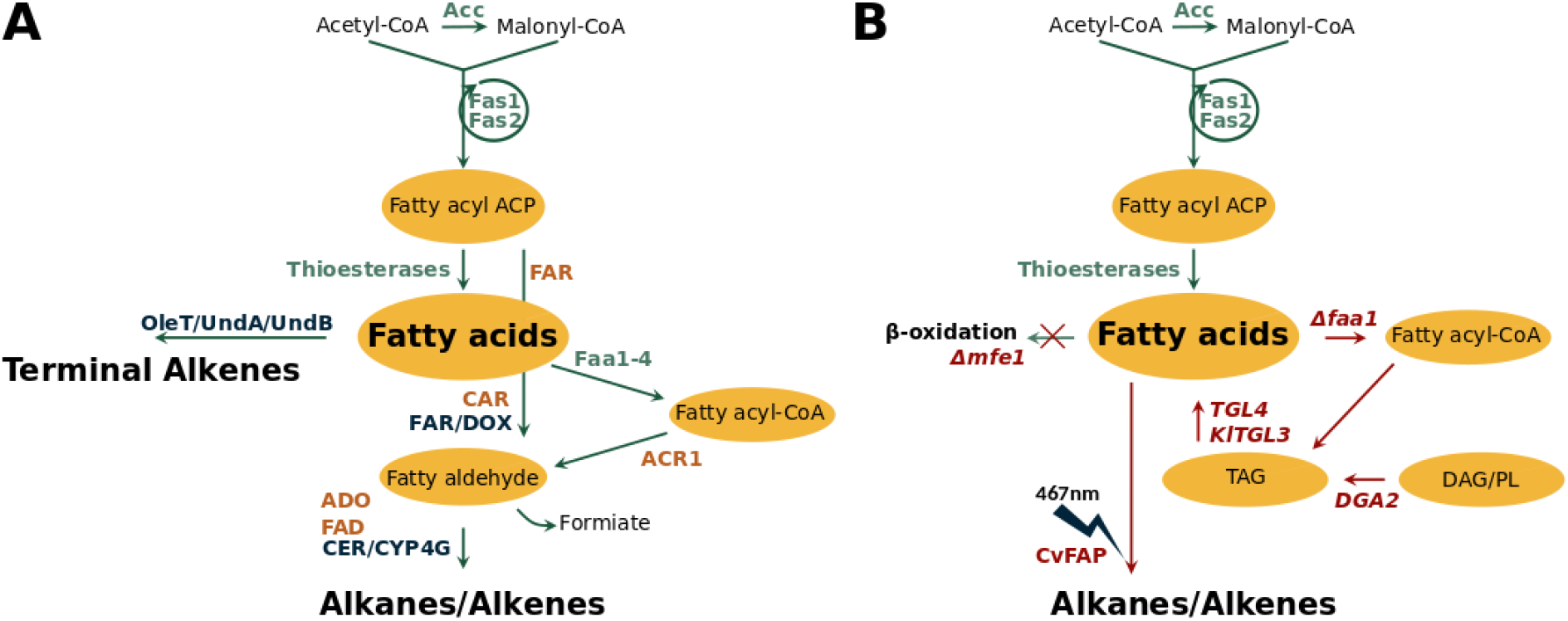
A: Previously described pathways for hydrocarbon production with yeast (modified from [3]). *Y. lipolytica* enzymes are shown in green, intracellular metabolites in black. Orange coloured enzymes are investigated in [3], dark blue coloured enzymes are reviewed in [35]. B: Expression of CvFAP, with modifications of strain JMY5749 are shown in red, characterized in this study.

In this work, we present the expression of CvFAP for the first time in an oleaginous yeast - *Yarrowia lipolytica*. Using different strain backgrounds, we detected up to 10.9 mg/L of hydrocarbon formation in a strain with an increased pool of free fatty acids (FFA) [5]. In order to increase these titers we engineered and 3D-printed labware to test different light regimes both in microwell plates and in repeated fed-batch processes in bioreactors. Additionally the influence of CvFAP mutant S121F was evaluated in bioprocesses. The highest titer of 58.7 mg/L was reached in a bioprocess feeding glycerol, using blue light with a wavelength of 465-470 nm and an continuous intensity of approx. 200 µmol quanta photons m^−2^ s^−1^.

## 2. Results and discussion

Due to low turnover number, the need of a coupled electron transfer system and the reactive and toxic substrate fatty aldehyde, the expression of ADO in yeast was connected with high efforts of metabolic engineering but small hydrocarbon yields (Tab.1). In contrast, the CvFAP enzyme directly utilizes FFA as its substrate as well as the readily available cofactor FAD. The catalysis is directly driven by the photons of blue light and hence tightly controllable. Unlike the AAR/ADO pathway, no additional genes for cofactor recycling are required [4].

### 2.1 Expression and characterization of CvFAP in *Yarrowia lipolytica* using YaliTAR

With regard to a rapid characterization, an *in vivo* DNA assembly strategy mediated by *Y. lipolytica* was carried out. In contrast to *S. cerevisiae*, which mainly employs homologous recombination as a DNA repair mechanism, in *Y. lipolytica* non-homologous end-joining (NHEJ) is preferred [6]. As a consequence, several DNA assembly methods developed for baker’s yeast are not directly transferable or applicable. In previous studies, efficient homologous recombination for genomic integration with short length flanking fragments was successfully shown for *Yarrowia Δku70* mutant strains [7]. In order to transfer the frequently used baker’s yeast transformation-associated recombination (TAR) [8] for assembling the centromeric CvFAP expression plasmid within *Yarrowia*, co-transformation of the backbone and corresponding insert in a *Δku70-*strain background (H222 SW1) was successfully performed. The insert contains a *Y. lipolytica* codon-optimized sequence of the CvFAP gene flanked in front by a *TEF1* promoter and a *XPR2* terminator at the end.

Positive constructs (verified by sequencing) were grown on lipid body formation inducing YSM medium, containing 5% D-glucose as carbon source, under exposure with a readily available plant LED-light for 96 h. Under these light conditions an intracellular titer of 112.1+/-31.4 µg/L of hydrocarbons could be detected. In a dark experimental setup 1.5+/-1 µg/L were detected. The empty vector control revealed no detectable production of hydrocarbons (Suppl., Tab. S1). The cloning method was coined as “YaliTAR”, derived from its *S. cerevisiae* analogon and enables direct characterization in *Y. lipolytica*, without the need for a shuttle host. The method can be generally applied for any other target gene and in particular deployed for rapid complementation of desired enzymatic activity.

### 2.2 Alkane production with CvFAP in different *Y. lipolytica* strain backgrounds

To evaluate the influence of different strain backgrounds with respect to fatty acid availability, we transformed the *C*vFAP expression vector into two different strains. We chose the laboratory strain H222 with knocked out beta-oxidation for increased lipid accumulation and deleted *ALK1* gene for inhibition of alkane degradation (S33001), as well as strain JMY5749 (genotype of JMY5749/CvFAP highlighted in Fig. 1B), an overproducer of free fatty acids (FFA), described in [5], for enhanced substrate availability. A blue light LED-strip with a more distinct wavelength range (465-470 nm) was used. The duration of cultivation was 96 h to impede complete depletion of glucose in order to inhibit alkane degradation through C-catabolite repression [9]. The cell dry weights of both strains at the end of the cultivation were in a similar range (3.6-4.4 mg/ml, Suppl., Tab S2). A nearly 30 times higher hydrocarbon titer was achieved for JMY5749- in comparison to the S33001-strain background under exposure of blue light (1.551 +/- 0.247 mg/L in contrast to 0.056 +/- 0.004 mg/L, Suppl., Tab S1). Despite reduction of alkane monooxygenase activity [10], as well as incapacity of fatty acid degradation, formation of hydrocarbons using S33001-strain background remain clearly lower than for JMY5749. In contrast, the latter features an increased lipase activity and thus provision of substrate in higher intracellular concentrations which underlines the requirement of the CvFAP for free fatty acids.

By optimizing the extraction method through usage of a ball mill, lowering sample volume and adaptation of GC method (see methods section), a titer of 10.87+/-1.11 mg/L total hydrocarbons could be detected, using JMY5749/CvFAP (shown in Fig. 2). Most of the hydrocarbons produced were heptadecane, 8-heptadecene and 6,9-heptadecadiene in similar levels followed by pentadecane and 7-pentadecene. Furthermore, the measurement of total fatty acids revealed a significant lower intracellular titer of 35 mg/g to 21 mg/g compared to the empty vector control (Suppl., Fig. S4). The hydrocarbon spectrum, listed above, also matches the findings of Sorigué *et al*. (2017) [4] with *E. coli* and Huijbers *et al.*, (2018) [11] *in vitro* experiments. Both groups recorded the highest enzyme activity for long-chained fatty acids (C16-C18), as well as a higher conversion rates for fully saturated substrates, comparing octadecanoic acid with unsaturated oleic or linoleic acid.

**Figure 2.**
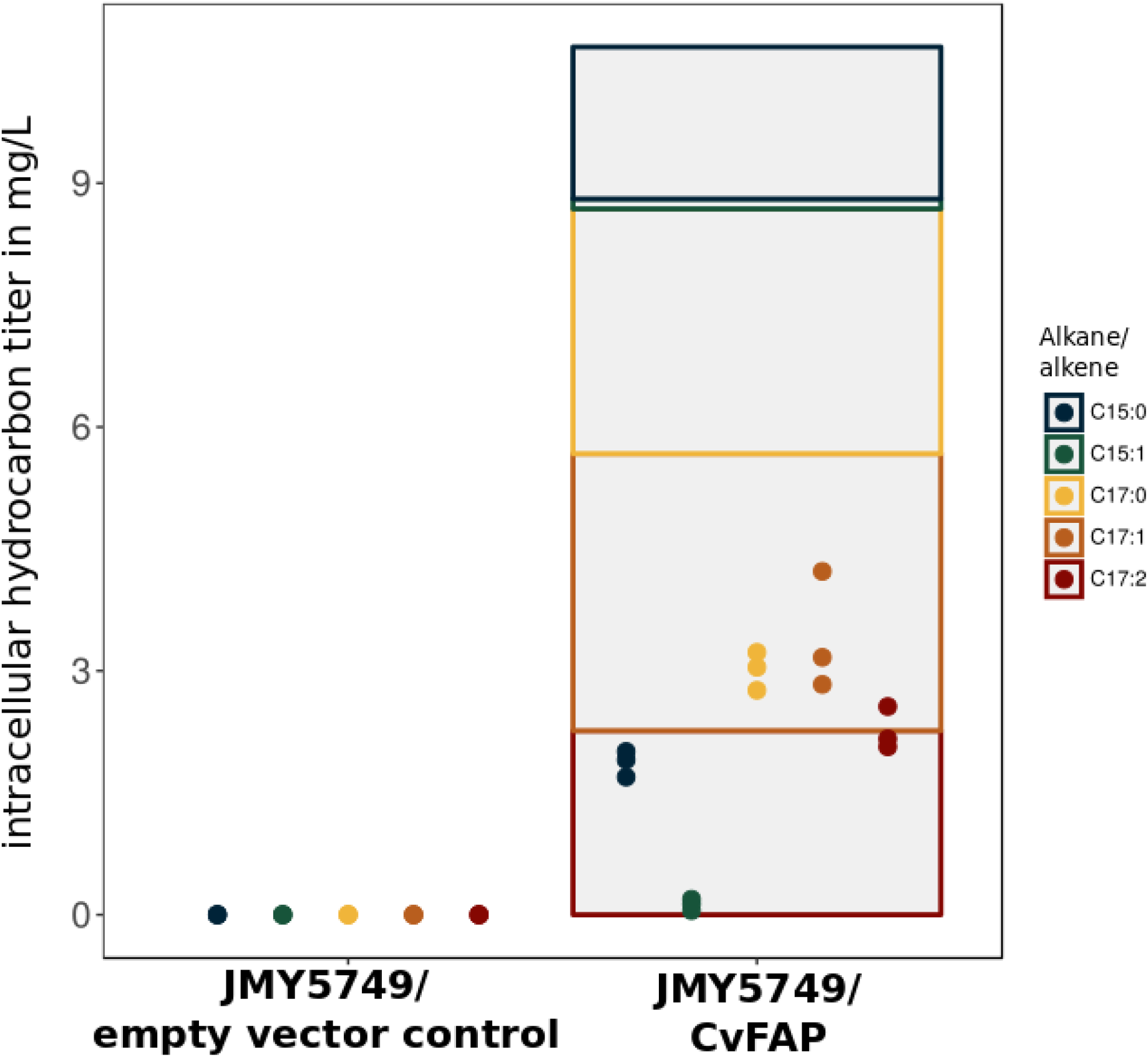
Alkane production with CvFAP expressed in *Yarrowia* JMY5749 in contrast to empty vector control (Strain background described in Tab. 2, medium composition in Tab. 3). The intracellular titer of each hydrocarbon is indicated by dots (in triplicates), the sum of hydrocarbons and the hydrocarbon composition is represented by a bar plot.

**Table 2.**
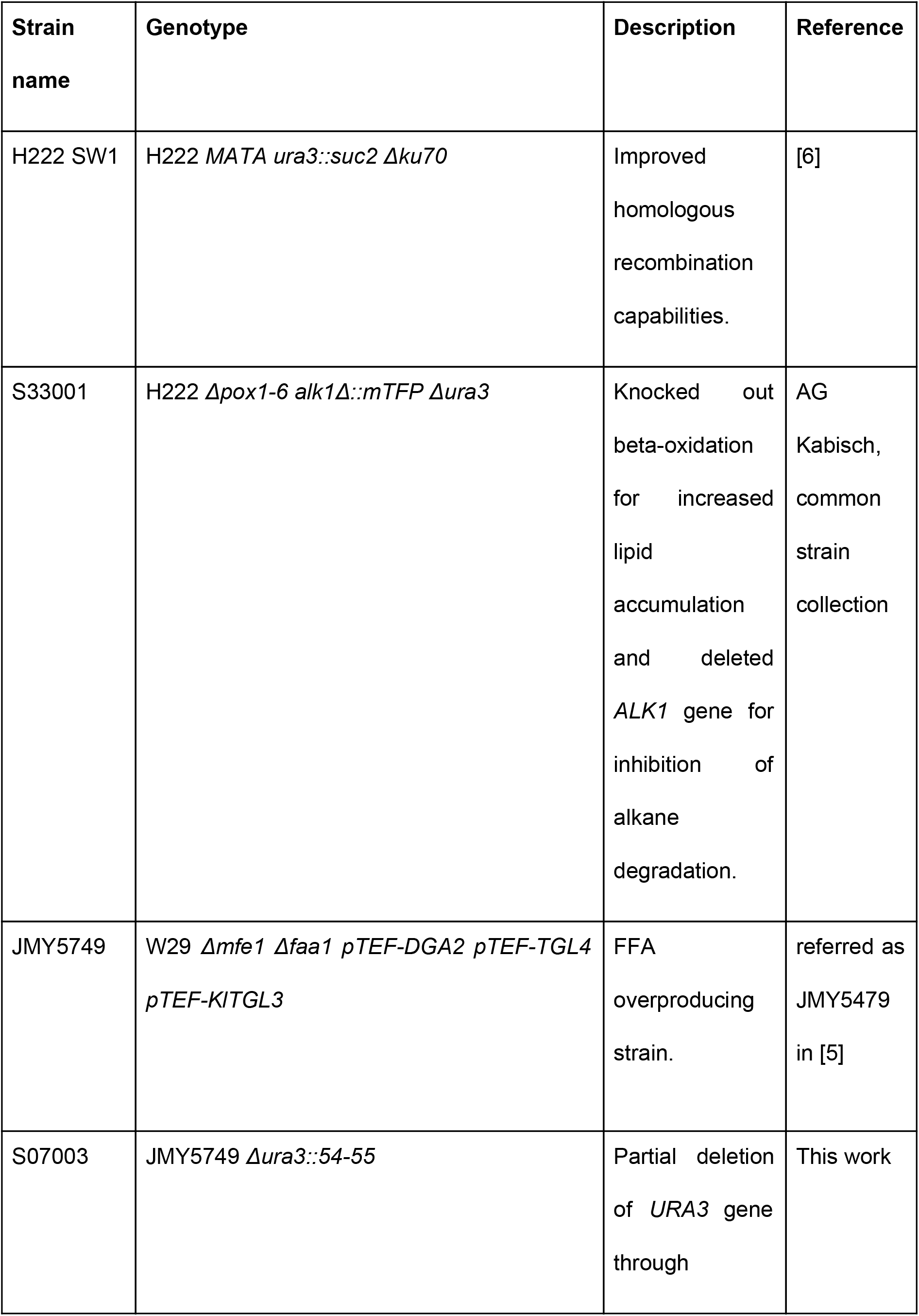
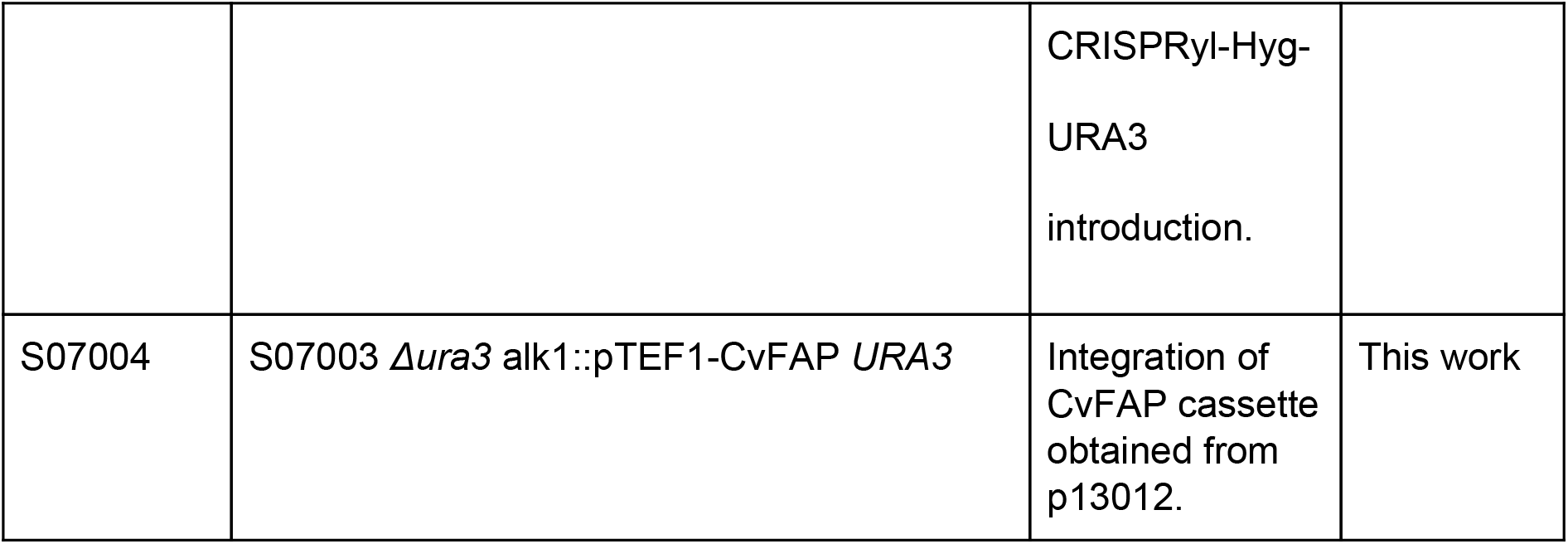
Yarrowia lipolytica strains used in this study

**Table 3.** *Yarrowia* mineral Salt Medium (YSM) for the induction of lipid droplet (LD) formation based on [33,34]. The medium was composed as a cost-effective alternative to common LD inducible media and designed for fed-batch cultivations with *Y. lipolytica*.

| Base compounds | Chemical formula | Conc. [g/L] |
| --- | --- | --- |
| Sodium phosphate dibasic | $\text{Na}_2\text{HPO}_4 \cdot 2\text{H}_2\text{O}$ | 1.6 |
| Potassium dihydrogen phosphate | $\text{KH}_2\text{PO}_4$ | 0.092 |
| Ammonium sulfate | $(\text{NH}_4)_2\text{SO}_4$ | 0.5 |
| **Further supplements** |  |  |
| Magnesium sulfate heptahydrate | $\text{MgSO}_4 \cdot 7\text{H}_2\text{O}$ | 0.7 |
| Calcium chloride hexahydrate | $\text{CaCl}_2 \cdot 6\text{H}_2\text{O}$ | 0.1 |
| yeast extract | - | 0.5 |
| D-glucose | - | 50 |
| **Trace elements** |  |  |
| Boric acid | $\text{H}_3\text{BO}_3$ | 0.0005 |
| Manganese sulfate monohydrate | $\text{MnSO}_4 \cdot \text{H}_2\text{O}$ | 0.0004 |
| Zinc sulfate heptahydrate | $\text{ZnSO}_4 \cdot 7\text{H}_2\text{O}$ | 0.0004 |
| Ferric chloride hexahydrate | $\text{FeCl}_3 \cdot 6\text{H}_2\text{O}$ | 0.006 |

The empty vector control did not show any hydrocarbon formation. No hydrocarbons (alkanes C8-C20) could be detected in the supernatants of any samples (data not shown).

### 2.3 Examination of process parameters and strain development using a custom-made device for cultivations in 24-well scale

In order to determine an appropriate light regime, comprising intensity, duration of exposure or effect of light pulsing, a medium throughput approach for light-dependent cultivation in 24-well plates was established. Besides the tracking of optical density, the cultivation volume of 750 µl was sufficient to allow endpoint measurement of intracellular hydrocarbons. Originating from a custom-made LED-device [12], adapters for LED-matrix and 24-well plates, as well as a universal plate-holder for incubators were fabricated (Fig. 3 AB). The rapid prototyping of custom labware proofed to be a very valuable tool in this work. Using free, open source software like openSCAD, readily available designs from previous publications and a 3D printer, we were able to parallelize the workflow using the 24-well LED plates in a shaker and avoid evaporation (data not shown) without resorting to expensive, commercial solutions.

**Figure 3.**
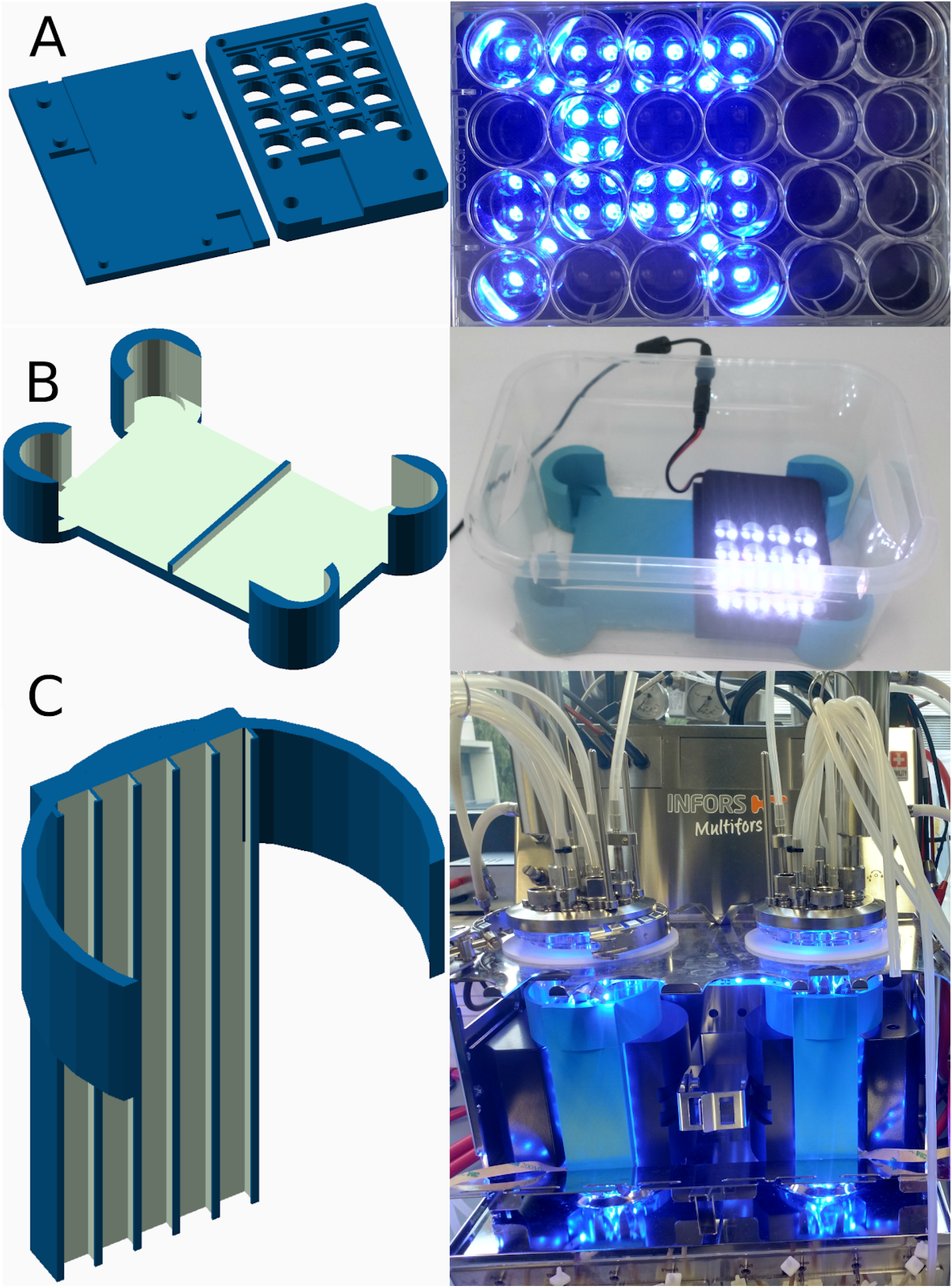
3D printed custom labware for evaluation of light regimes. A: Rendering and picture of a LED-matrix plate for testing light regimes in microwell plates. B: Low evaporation setup with custom microwell plate-holder and LED-matrix in a low cost plastic box. C: Rendering and picture of LED-strip-holder attached to bioreactors.

Cultivation of JMY5749 carrying a CvFAP-plasmid at high cell densities on glucose-containing YSM medium showed the highest intracellular hydrocarbon titer using maximum LED intensity of 29-32 µmol quanta m^−2^ s^−1^ per well and continuous lightning. Short light pulses with breaks of 100 ms or 5000 ms, as well as an intensity reduced by half, led to significant reduced hydrocarbon formation (Fig. 4 A black dots, Anova1 in Suppl.). The growth, determined by optical density, was not affected by any of the lighting conditions (Suppl., Fig. S5 A). To further investigate a putative low impact of continuous lighting or even pulsing, the measurements were repeated by adjusting the initial high OD_600_ to 0.1 (Fig. 4 A, orange dots). Equally, a reduction of the growth rates at given intensities could not be detected (Suppl., Fig. S5 B). In contrast to the first approach, the lighting with half intensity led to a similar hydrocarbon titer, which indicates the importance of fine-tuning light exposure and cell growth (Anova2 in Suppl.). To examine further genetic contexts, strains harboring a genomic integration of *Cv*FAP were characterized with the help of the 24-well LED-device. By applying continuous lighting at highest possible intensity without pulsing, all clones showed similar hydrocarbon titer after 96 h of cultivation in YSM medium (Supp., Fig. S5 C). Due to differences in alkane/alkene composition (Fig. 4 B), strain S07004 EK1, showing a mutated CvFAP sequence (S121F in WT, S61F for truncated CvFAP without signal peptide; Suppl., Seq 1; rendering based on the wild type and the mutant protein structure is shown in Supp., Fig. S6.) with confirmed integration in *ALK1* locus, as well as strain S07004 with correctly integrated *Cv*FAP wild type sequence, were chosen for a more precise characterization in half-automated, repeated fed-batch bioreactor cultivations.

**Figure 4.**
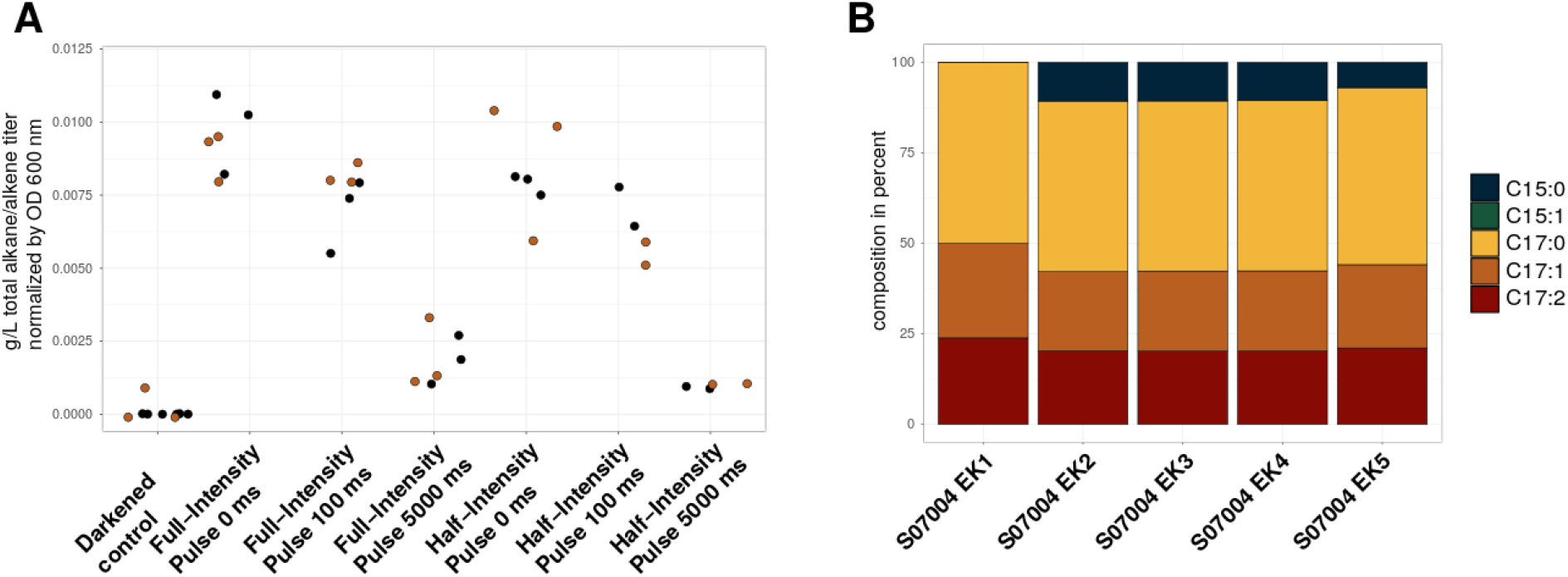
A: Endpoint measurements of total hydrocarbons formed in microwells with different light regimes. Inoculation with high (set OD_600_ of 10; black dots) and low (OD_600_ was set to 0.1; orange dots) initial cell densities. Full light intensity was determined as 28.7-32.3 µmol quanta m^−2^ s^−1^ per well. Light regimes were tested in triplicates, except for half intensity, pulse 100 and 5000 ms, which were cultivated in duplicates. Measurements are coloured in black. B: Alkane/alkene compositions of endpoint measurements for cultivations (initial OD_600_ 0.1) of different S07004 clones, harboring the genomic integration of *Cv*FAP cassette. Strains were cultivated in triplicates, expect for n=2 S07004 EK4.

### 2.4 Characterization of different strains and light intensities, using bench-scale bioreactors equipped with custom-made holders for LED-stripes

To perform light-dependent bioreactor cultivations, custom-made holders for the previous utilized LED-stripes were fabricated using a 3D printer for the reactor vessels (Fig. 3 C) of a common bench-scale bioreactor. The construction of the holding clips attaching the LED-strips to the bioreactors ensured reproducible lighting conditions (487-560 quanta photons m^−2^ s^−1^ for full light intensity). The intensity of the LED-stripes was controlled by a standard laboratory power supply. The ratio of single LED-light intensity and current was determined (Suppl., Tab. S3) in order to facilitate reproducible blue light intensities. Batch medium contained 30 g/L carbon source and 5 g/L ammonium sulfate for the generation of biomass. After batch phase, a pulse of 30 g/L C-source containing feed solution was added when C-source was depleted (detected by the rise of dissolved oxygen (DO)). By omitting a nitrogen source in the feed medium, an increased C/N ratio and thus an increased formation of free fatty acids should be achieved [5]. The compositions of batch and feed medium, as well as a detailed description of the bioreactor and cultivation conditions are listed in the material and methods section. The sequence for DO-dependent automated feeding can be found in the Supplementary file.

In the light of above-described 24-well cultivations, clones S07004 EK1 (CvFAP S121F) and EK4 (CvFAP WT) were cultivated in triplicates. In contrast to previous experiments, glycerol was selected as carbon source, in particular due to its availability as a sidestream in biodiesel production. While growth parameters like DO concentration and feeding intervals were similar (Fig. 5 A), minor differences in intracellular total hydrocarbon titer and cell dry weight could be detected (Fig. 5 B,C). The highest titer of 16.71 mg/L was reached by S07004 EK1 (CvFAP S121F), furthermore slightly higher biomass was obtained. In contrast to the hydrocarbon composition obtained in the 24-well plates, pentadecane could be detected after 15 h of cultivations in all bioprocesses of both, wild type and the S121F variant. These differences between the bioreactor and 24-well plates could be caused by a three-fold increased light intensity (max. 60-90 µm quanta m^−2^ s^−1^) in the bioreactor underlining that while the microwell cultivations can be used as a first screening a controlled process environment is required for in depth analysis.

**Figure 5.**
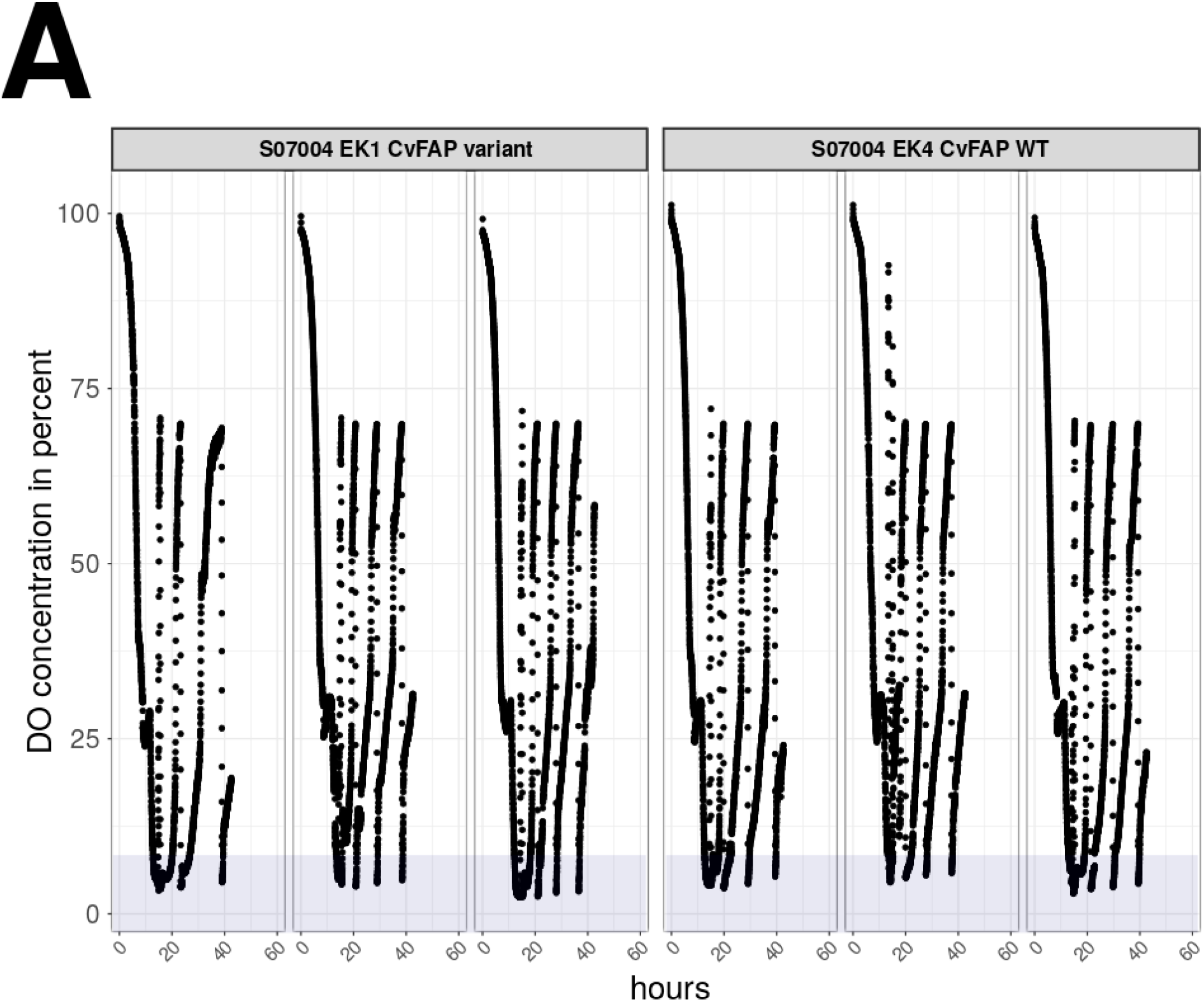
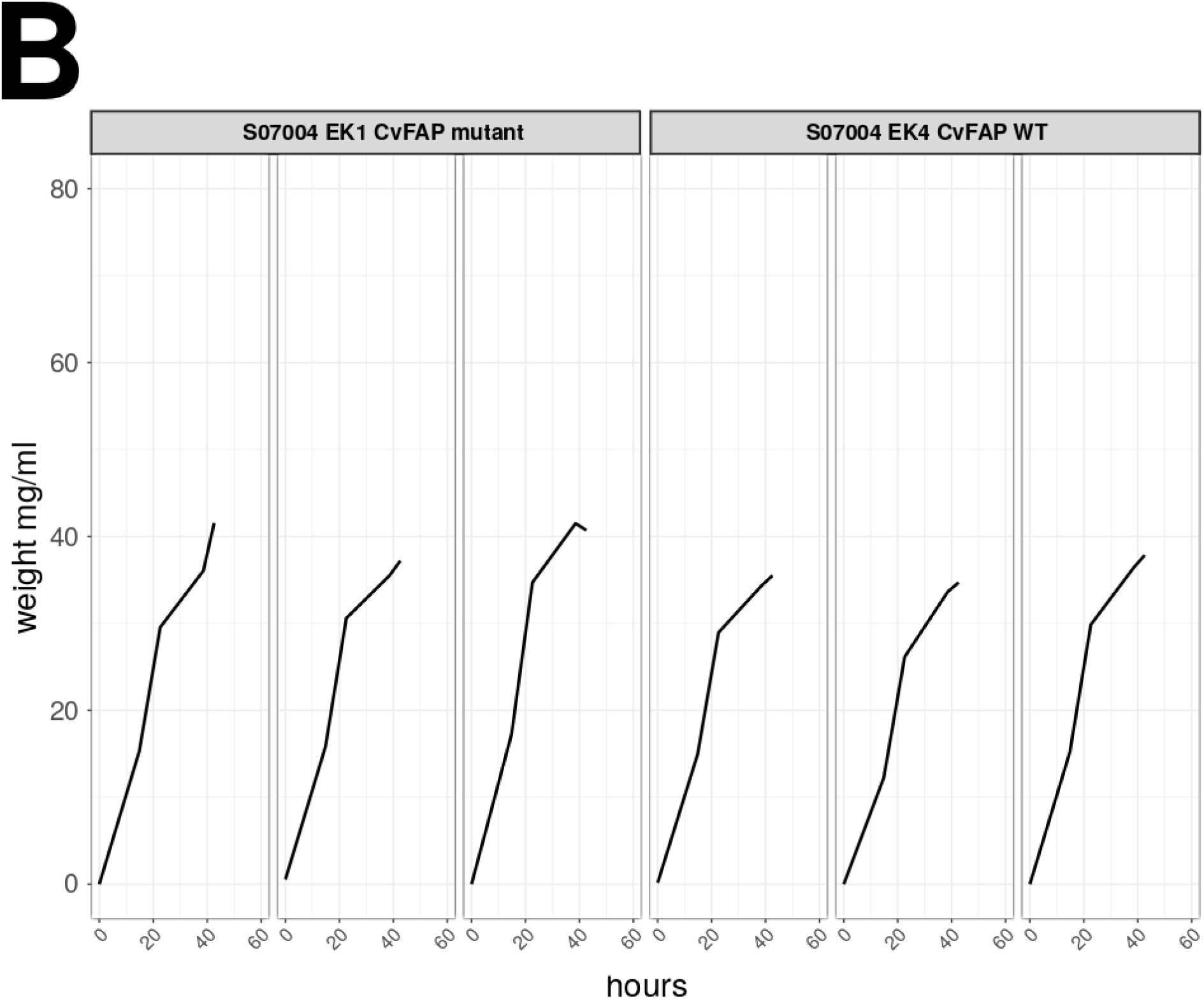
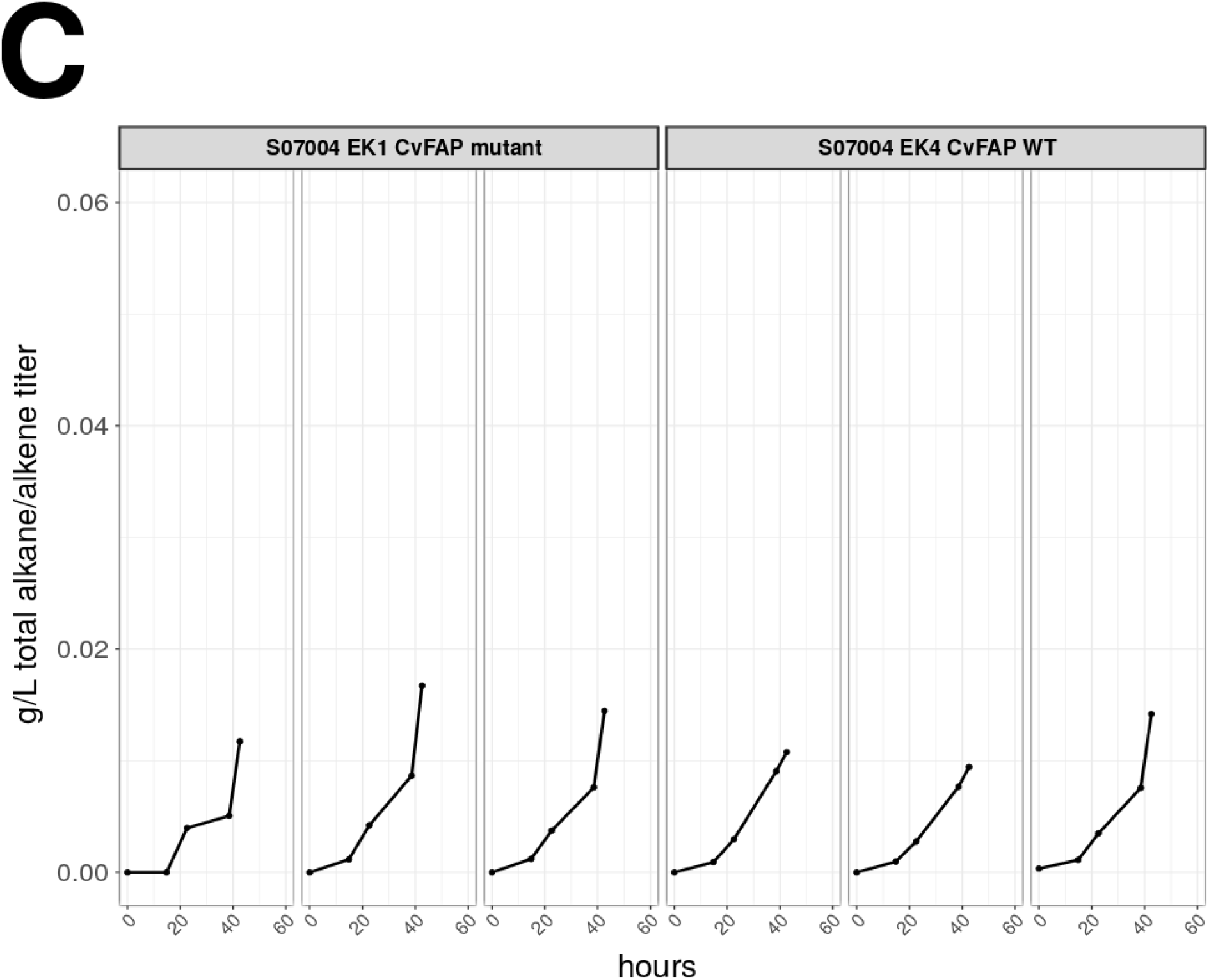
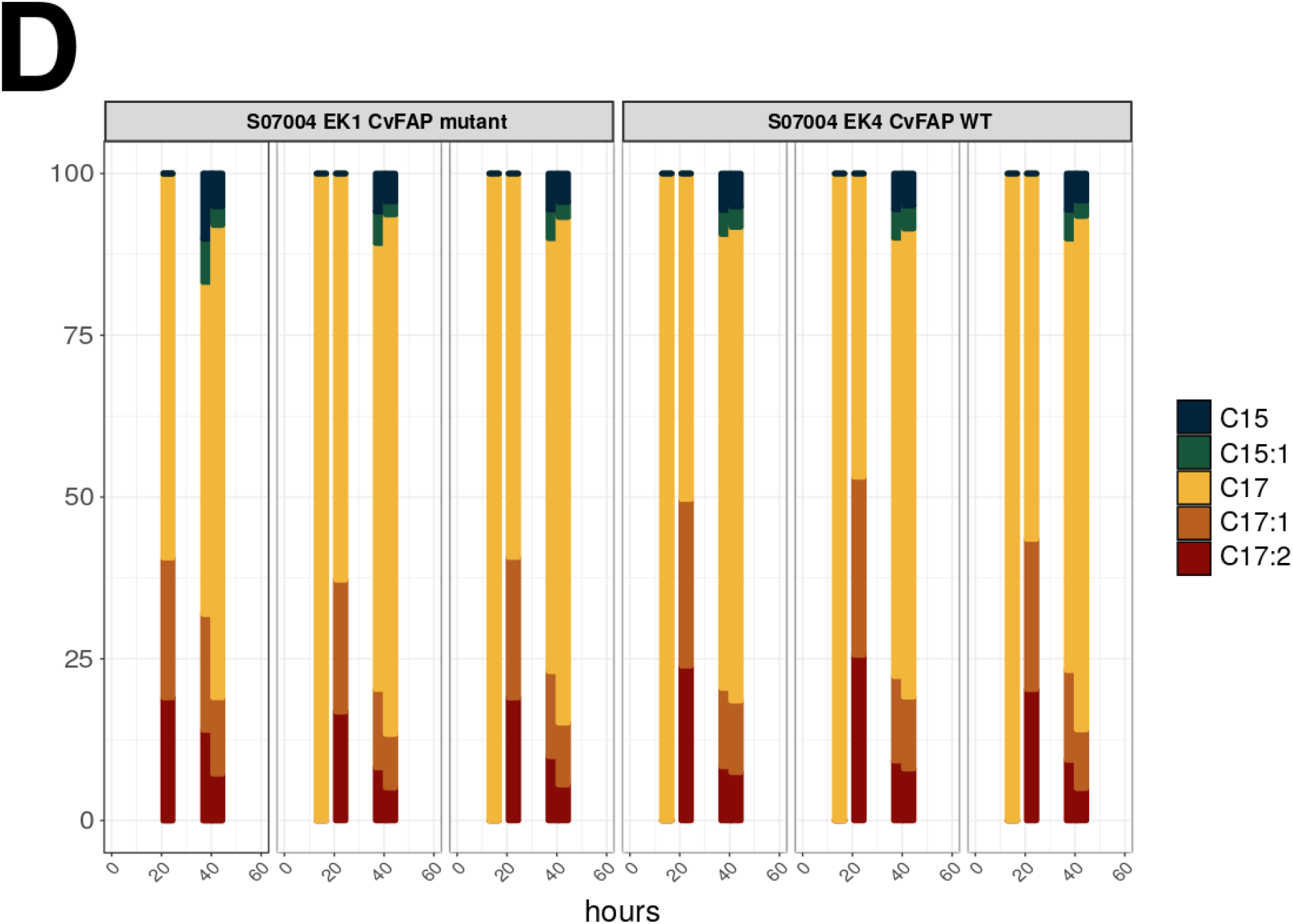
Comparison of CvFap variant (S07004 EK1; S121F) and wild type (S07004 EK4), cultivated in triplicates. A: DO concentration as a representation of metabolic activity. Light intensity (height) and exposure time (width) are indicated by the blue shaded areas (e.g. the light intensity was set to ∼60 µm quanta m^−2^ s^−1^ which is ∼1/10th of full intensity) B: Cell dry weight (cdw), to determine biomass accumulation in mg/ml. C: Total intracellular hydrocarbon titer in g/L, concentrations in relation to cdw are shown in the Suppl., S9 C. D: Distribution of measured hydrocarbons in percent.

Despite a higher titer of 8-heptadecene and 6,9-heptadecadiene formed by the CvFAP WT at 20 hours the composition of the remaining hydrocarbons showed a high similarity between the WT and the CvFAP variant (Fig. 5 D). A better performance of S07004 EK1 could also be verified by supplying glucose as carbon source in an additional experiment (Suppl., Fig. S7 ABC). When feeding glucose as C-source we noticed a less and less responsive DO-signal (Suppl., Fig. S7 A3), which we already knew from other experiments with *Y. lipolytica* (data not shown) and which resulted in a less robust performance of the feeding sequence used for automatic control. This effect does not occur when feeding glycerol (Suppl., Fig. S7 A). Therefore further experiments were performed using glycerol as C-source. Based on the better performance of S07004 EK1 further studies were continued with this strain harboring the amino acid exchange. A structural analysis based on the published structure of this CvFAP variant revealed a minimum distance of 12 Å, between the phenylalanine ring and flavin adenine dinucleotide, but was shielded by secondary structures (Suppl., Fig S6). According to literature, a different positioning of the functional carboxyl group to the cofactor could have a strong influence on substrates conversion rates or yields [11]. Thus, an indirect influence due to altered coordination of intermediate AA residues should be examined in future studies.

### 2.4.1 Improvement of hydrocarbon production depends on orchestration of light intensity and growth

In the 24-well experiments no reduction in growth could be detected by using a maximum intensity of 32 µmol quanta m^−2^ s^−1^ per well. The LED-strips attached to the bioreactores allowed for a seventeen times higher light intensity (approx. 560 µmol quanta m^−2^ s^−1^). In order to obtain the highest amount of total hydrocarbons, four different light settings were evaluated. Besides full intensity and no blue light control (ambient light), also half intensity (approx. 200 µmol quanta m^−2^ s^−1^) and an induction with full intensity, 16 h after inoculation were tested (Fig 6, Suppl., Fig. S8). As an online measurement of the metabolic activity of the cells we used the values of the DO-probe (Fig. 6 A). Full intensity led to a decreased amount of feeding cycles and a longer interval between each feeding pulse in comparison to the no-light, half intensity and late-light-induction processes (Fig. 6 A). In contrast to the other conditions and despite similar levels of biomass formation, intracellular octadecanoic acid (free fatty acid or incorporated in triacylglycerides) concentration were relatively elevated (Suppl., Fig. S8 A) during full light conditions. Furthermore, the reduced feeding was accompanied by lower formation of extracellular metabolites like citrate and polyols (Suppl., Fig. S8 D). Interestingly, the highest total hydrocarbon formation with regard to the cell dry weight, 2.5 % (7.12 mg/L), could be obtained after 20 h of cultivation (Suppl., Fig. S8 C). On average, the remaining light-dependent bioprocesses revealed a maximum total hydrocarbon formation of 0.14 % cdw (half of intensity: 0.169 %, 0.151 %, 0.168 %; late induction: 0.081 %, 0.158 %, 0.111 %), primarily owed to the more rapid increase of the biomass (Fig. 6 B). The severe growth impairment with full light conditions from the onset of the bioprocess, suggest that the CvFAP sequesters most fatty acids formed. After almost 20 hours of lag-phase the cells seem to recover and resume growth but stop producing hydrocarbons. In literature high light intensity, especially irradiation in the range of 450 nm, is linked to a phenomenon called photoinactivation, which results in *S. cerevisiae* cells which are in a viable but nonculturable state. As photosensitizers, flavins and porphyrins are discussed [13]. Furthermore, especially for blue light, a significant effect on yeasts respiratory oszillation is described [14]. To what degree these findings are transferable to *Y. lipolytica* needs to be examined in future studies.

**Figure 6.**
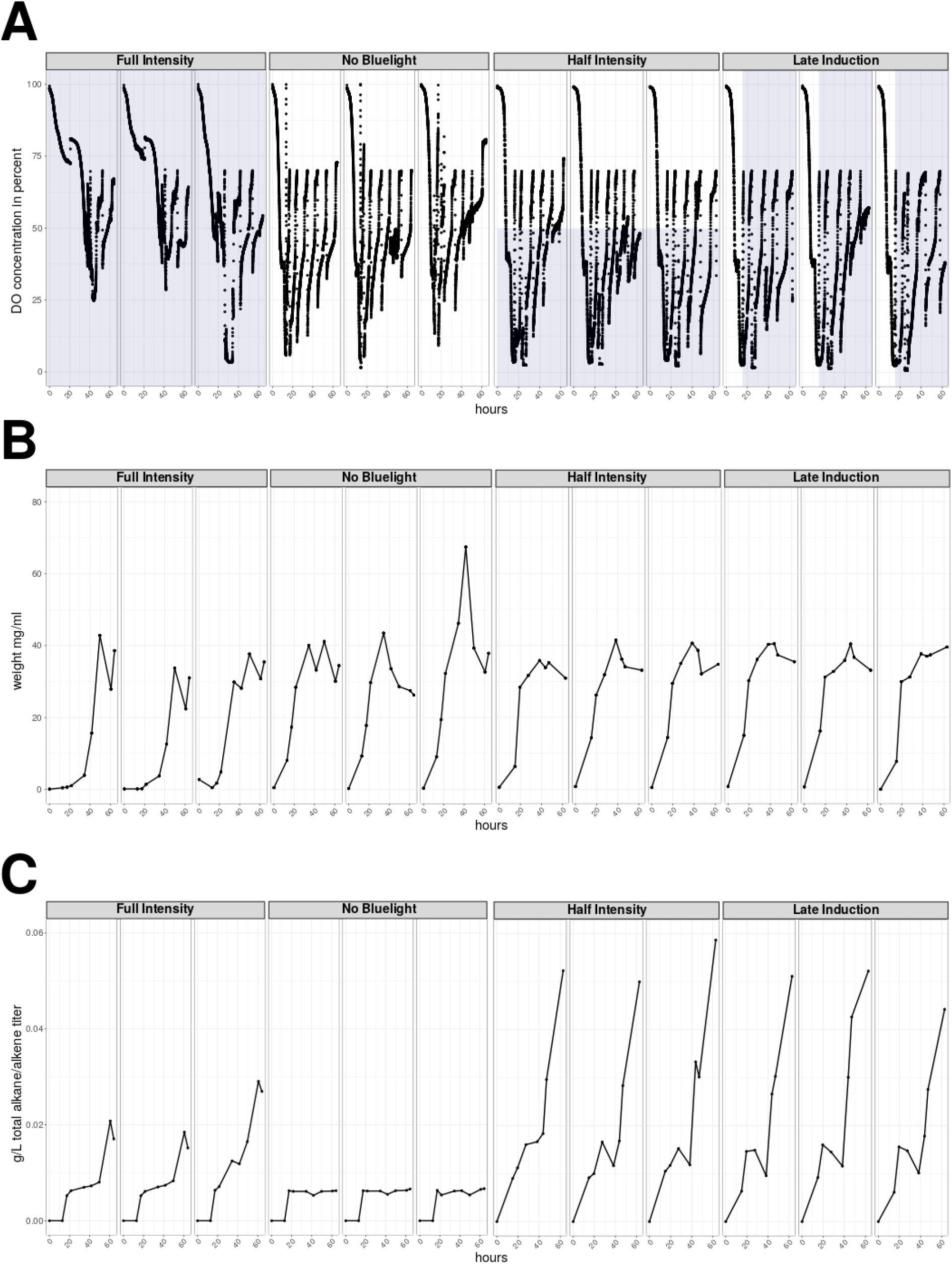
Bioprocesses with four different light regimes were characterized in triplicates by cultivation of strain S07004 EK1. Light intensity (height) and exposure time (width) are indicated the blue shaded areas. For full intensity, the light intensity was set to 545 µm quanta m^−2^ s^−1^, while half intensity reached 250 µm quanta m^−2^ s^−1^. For the no blue light control, fermenter vessel were shielded from neighboring blue light but were still affected by ambient light. For the late induction experiment, full light intensity was switched on 16h after inoculation. A: DO concentration in percentage. Light intensity and exposure time are indicated by blue coloured areas.

Considering the absolute total hydrocarbon formation, best results were achieved using half of intensity with a maximum of 58.69 mg/L, closely followed by late induction experiments (52.23 mg/L). To our knowledge and in comparison to other studies harnessing *Y. lipolytica* as host organism, these are the highest hydrocarbon titer described so far. Taken in account glycerol as a C-source for production of hydrocarbons, it is also the highest titer using yeasts. Expression of a heterologous α-dioxygenase from rice and a cyanobacterial aldehyde-deformylating oxygenase in *S.cerevisiae* in combination with feeding 10 mg/L fatty acids as precursors using a buffered medium resulted in slightly higher formation of 73.5 mg/L long-chained hydrocarbons. Higher concentrations were also reached by implementing the AAR/ADO pathway into *E. coli* (see Table 1 for comparison with recent studies).

**Table 1.**
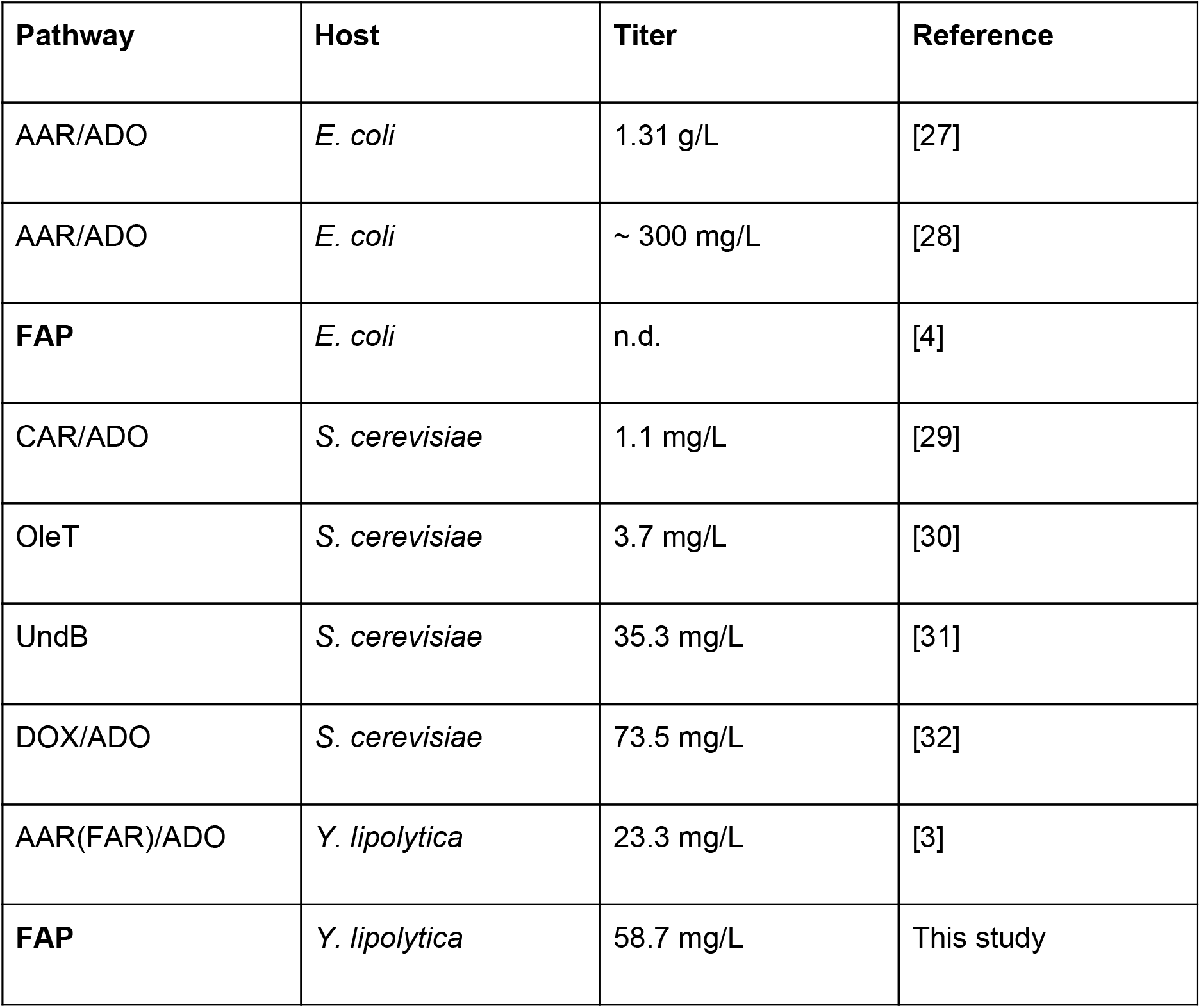
Hydrocarbons produced by selected organisms, expressing different heterologous pathways. The enzyme abbreviations are namely AAR for acyl-ACP reductase, ADO for aldehyde-deformylating oxygenase, FAP for fatty acid photodecarboxylase, OleT for 450 fatty acid peroxygenase/decarboxylase, UndB for desaturase-like enzyme, DOX for α-dioxygenase.

| Pathway | Host | Titer | Reference |
| --- | --- | --- | --- |
| AAR/ADO | *E. coli* | 1.31 g/L | [27] |
| AAR/ADO | *E. coli* | ~ 300 mg/L | [28] |
| **FAP** | *E. coli* | n.d. | [4] |
| CAR/ADO | *S. cerevisiae* | 1.1 mg/L | [29] |
| OleT | *S. cerevisiae* | 3.7 mg/L | [30] |
| UndB | *S. cerevisiae* | 35.3 mg/L | [31] |
| DOX/ADO | *S. cerevisiae* | 73.5 mg/L | [32] |
| AAR(FAR)/ADO | *Y. lipolytica* | 23.3 mg/L | [3] |
| **FAP** | *Y. lipolytica* | 58.7 mg/L | This study |

A decrease in hydrocarbon production in the form of stagnating or declining values was observed within all light processes. This indicates a putative degradation of formed alkanes or alkenes. In *Yarrowia lipolytica*, presence of *n*-alkanes lead to transcriptional activation of alkane-degrading enzymes. While the main monooxygenase (*ALK1*) responsible for hydrocarbon degradation is deleted in the strains used in this study, the remaining ALK2-12 enzymes are sufficient for the degradation of long-chain hydrocarbons [15]. Nevertheless the corresponding genes are subject to transcriptional repression due to glycerol feeding [16]. For half intensity and late induction settings, a major decrease could be detected after 40 h of cultivation. This coincides with a temporary glycerol exhaustion due to the broad settings of saturated oxygen limits of the feeding sequence (<40%, >70%; shown in Suppl., compare with extracellular glycerol titers, shown in Suppl., Fig. S8 D). Consequently, a tighter process control, taking into account the disadvantage of adaptation for specific strain backgrounds and light regimes, as well as further deletions of genes encoding alkane-degrading enzymes are likely candidates for further improvement of the hydrocarbon titer.

Generally, in all light-induced bioprocesses, with exception of the no blue light control, a predominant formation of heptadecane (C17:0) occurred (Suppl., Fig. S8 B). The effect was more pronounced when using glycerol instead of glucose as C-source (Suppl., Fig. S7 C). In no blue light control, similar amounts of C17:0 and unsaturated 8-heptadecene (C17:1) as well as 6,9-heptadecadiene (C17:2) were detected. Considering the fatty acid composition, this stands in contrast to the predominance of unsaturated fatty acids such as for example oleic acid (C18:1) over octadecanoic acid (C18:0). Thus a preference of the CvFAP in *Y. lipolytica* for saturated fatty acids can be assumed. This holds true for all detected fatty acids as shown in Fig. 7 and more detailed in Suppl., Fig. S9. While the lowest intracellular fatty acid titers could be assigned to hexadecanoic acid (C16:0), converted pentadecane (C15:0) partially showed third-highest titers of detected hydrocarbons. In contrast, relations of oleic acid (C18:1) and linoleic acid (C18:2) and the formed hydrocarbons 8-heptadecene (C17:1) as well as 6,9-heptadecadiene (C17:2). Values of palmitoleic acid in comparison to 7-pentadecene (C16:1, C15:1) imply lowest turnover. In the no blue light control these effects were not fully confirmed (Suppl., Fig. S9). The preference for saturated fatty acid could be explained with the findings for purified CvFAP enzyme which exhibits higher conversion rates for saturated fatty acids [11].

**Figure 7.**
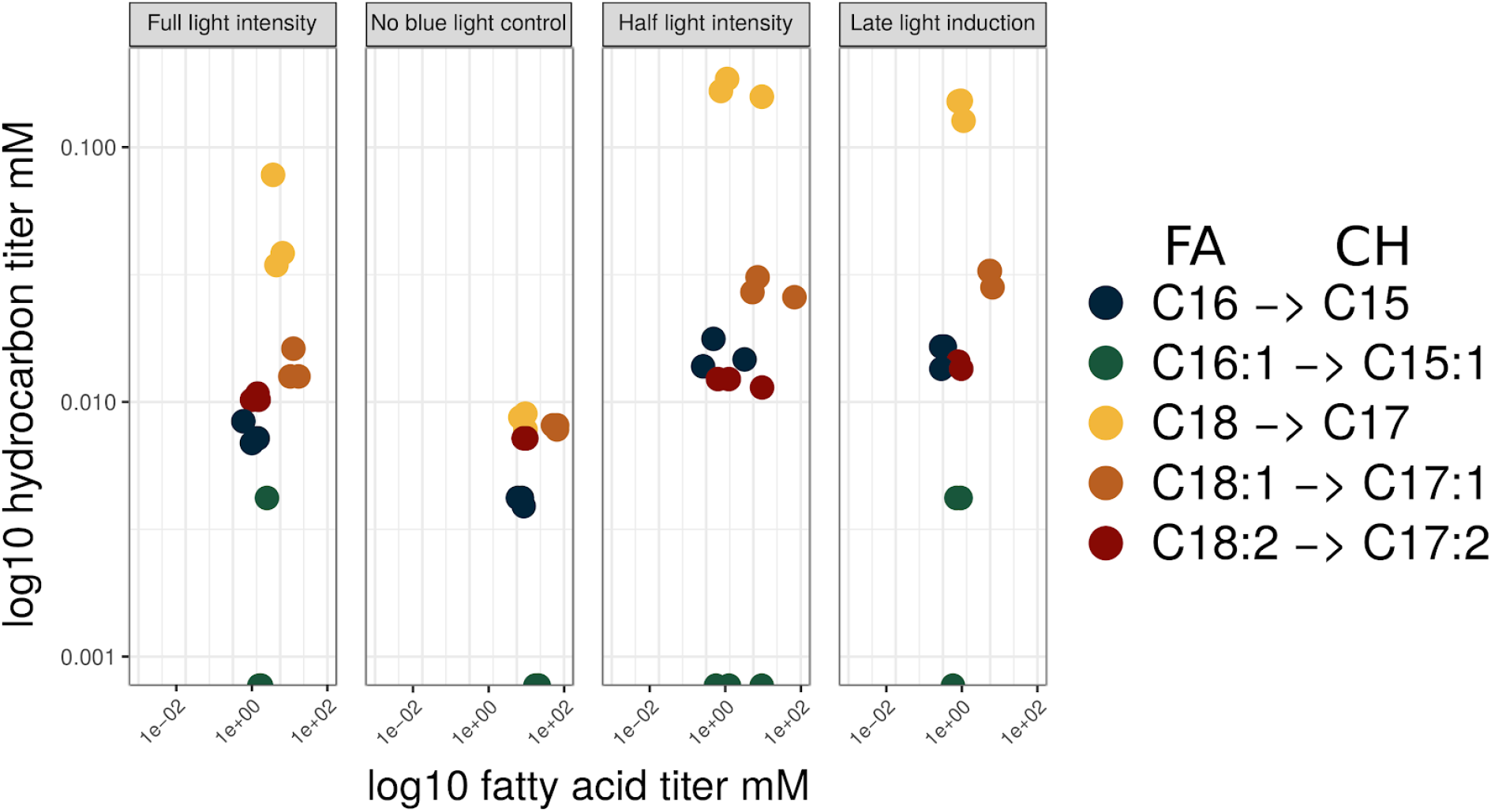
Amounts of fatty acids and hydrocarbons formed until the end of the cultivation. More detailed Figure in Suppl. Fig. S9.

## 3. Conclusions

Expression of CvFAP in oleaginous, fatty acid secreting *Yarrowia lipolytica* under blue light exposure leads to the production of odd-numbered alkanes and alkenes with a predominant length of 17 and 15 carbons. Especially the absence of reliable and readily available, inducible promoters for *Y. lipolytica* makes this light-driven reaction desirable with respect to process control. For example by omitting the inducing wavelength, the enzyme could be produced from a constitutive promoter and catalysis was switched on only after sufficient amounts of fatty acids have accumulated. 3D printing and readily available LED technology are especially interesting technologies to combine with light-driven bioprocesses, enabling researcher to rapidly develop custom labware.

Future strain engineering should include aspects such as increasing gene copy numbers, reduce secreted metabolites and modify fatty acid profiles. Process should take CO_2_ released during decarboxylation into consideration and could include recently described carbon dioxide (CO_2_) fixation approaches [17] [18].

## 4. Material and Methods

### 4.1 Transformation-associated recombination assisted by *Y. lipolytica* (YaliTAR) for rapid construction of simple replicative vectors

For time-saving *in vivo* assembly, Yarrowia strain H222 *Δku70* was co-transformed [19] with linearized replicative vector p15018 backbone (Suppl., Fig. S1 A), previously digested by MluI and NotI, as well as codon-optimized CvFAP fragment (Suppl., Seq. 1), including 43 bp homologous sequences to *TEF1* promoter (additional 6 bp MluI restriction site for further promoter exchanges) and *XPR2* terminator of p15018. CvFAP fragment was purchased at Baseclear B.V., without the predicted targeting sequence as shown in [4]. Oligonucleotides for amplification of overlapping fragments are listed in Suppl., Tab. S4. Positive clones were selected on YPD2% agar plates including 400 µg/ml hygromycin after 1-2 days of incubation at 30 °C. Vectors, recovered from 4 out of 14 colonies, were verified by sequencing, whereby 50 % showed the correct sequence (named p13001, but “CvFAP” in figures). YaliTAR method was also applied for the exchange of a marker gene of the Cas9 expression vector pCRISPRyl [20,21], which was provided by Ian Wheeldon (Addgene plasmid # 70007). Exchange from leucine to hygromycin marker resulted in vector p55001, verified by sequencing. Further integration of sgRNA was performed using SLiCE *in vitro* method, described below (Oligos listed in Suppl., Tab. S4, vectors in Tab. S5). *Yarrowia* strains H222 *Δku70*, S33001 and JMY5749 (Tab. 2) were transformed with p13001 vector.

### 4.2 *Y. lipolytica* strain construction

Backbone, originating from integrative vector p33001 (Suppl., Fig. S1 B), and CvFAP cassette from previous described p13001 were amplified with overlapping homologous overhangs (Suppl., Tab. S4/S5). For the assembly of both parts, SLiCE method were used, described by [22] including minor deviations listed at [23]. This resulted in vector p13012. CRISPRyl-Hyg-URA3 (p94001) was constructed likewise, sgRNA was designed by CHOPCHOP v2 online tool [24][20]. Before successful integration in JMY5749 background, first deletion of *URA3* by transformation with p94001 and counter selection with 5-fluoroorotic acid [25] caused the required auxotrophy. Partial deletion of *URA3* in JMY5749 was verified by sequencing. To verify subsequent integration of CvFAP in *ALK1* locus, transformed cells were selected by growth on YNB ura^−^ plates. Positive clones were picked and checked by sequencing.

### 4.3 Cultivation conditions and sampling for shake flask experiments

5 ml YPD2% (referred elsewhere) for inoculation and 25 ml YSM medium (low mineral salt medium, according to Tab. 3) for the induction of lipid body formation were used.

Cultivations in shake flasks were performed at RT (H222 *Δku70*/CvFAP and empty vector control) or 28°C and 180 rpm. The shakers were darkened as indicated. Light for the photoenzyme was provided by a commercial blue light LED-strip with an advertised wavelength of 465-470 nm (Fig. S2) or a common LED plant breeding light from Florally Inc. (Shenzhen, Guangdong, 518000,CN). Samples were taken after 96 hours for the determination of cell dry weight, intracellular and extracellular hydrocarbons and metabolites in the supernatant. For analytics, the entire volume of supernatant and cells were utilized (for improved extraction method, 1 ml of sample volume were chosen).

### 4.4 Cultivation conditions and sampling in a custom-made 24-well device

For testing the influence of different intensities and pulsing of the blue light required for photodecarboxylation a LED-matrix plate and a holder were fabricated using 3D printing. The setup is depicted in Suppl. Fig. S5, design and printing is described in material and methods section 6.9. For incubation in a shaker, a common plastic box with dimensions width/depth/height of 12.1/25.5/13.6 cm (RegaLux clear Box XXS, BAUHAUS), as well as 24-well sensor plates with glass bottom and darkened walls (Sensoplate glass bottom, Black, Greiner BIO-ONE, Austria) were used. Cells were grown on 750 µl YSM medium, according to Tab. 3, at 28°C and 180 rpm. For endpoint measurements, residual volume of cultivation broth were measured and used for hydrocarbon analytics, described below.

### 4.5 Bioreactor cultivations and sampling

Batch medium for fermentation contained 30 g/L carbon source (glucose or glycerol), 0.5 g/L yeast extract, 1.1 g/L MgSO_4_*7H_2_O, 0.2 g/L CaCl_2_*6H_2_O, 0.5 g/L MgCl_2_*6H_2_O, 0.075 g/L myo-Inositol, 1.36 g/L KH_2_PO_4_, 1.74 g/L K_2_HPO_4_, 0.2 mg/L CuSO_4_*5H_2_O, 1 mg/L FeSO_4_*7H_2_O, 0.2 mg/L MnCl_2_*4H_2_O, 0.2 mg/L Na_2_MoO_4_*2H_2_O, 0.2 mg/L ZnSO_4_*7H_2_O, 5 mg/L biotin, 100 mg/L D-pantothenic acid hemicalcium salt, 20 mg/L nicotinic acid, 60.8 mg/L pyridoxine hydrochloride, 20 mg/L thiamine hydrochloride and 5 g/L NH_4_Cl. Feed medium for fermentation contained 400 g/L carbon source (glucose or glycerol), 3.3 g/L MgSO_4_*7H_2_O, 0.6 g/L CaCl_2_*6H_2_O, 1.5 g/L MgCl_2_*6H_2_O, 0.45 g/L meso-Inositol, 2.72 g/L KH_2_PO_4_, 3.48 g/L K_2_HPO_4_, 0.6 mg/L CuSO_4_*5H_2_O, 3 mg/L FeSO_4_*7H_2_O, 0.6 mg/L MnCl_2_*4H_2_O, 0.6 mg/L Na_2_MoO_4_*2H_2_O, 0.6 mg/L ZnSO_4_*7H_2_O, 15 mg/L biotin, 300 mg/L D-pantothenic acid hemicalcium salt, 60 mg/L nicotinic acid, 182.4 mg/L pyridoxine hydrochloride, 60 mg/L thiamine hydrochloride and 0.01 g/L FeCl_3_*6H_2_O. Bioreactors (Infors multifors 2) were inoculated to an optical density at 600 nm of 0.1 in 300 mL of batch medium from overnight shake flask. Initial process parameters were pH 6.0, temperature of 30 °C, aeration at 1 lpm air, and agitation at 400 rpm. Afterwards the pH was adjusted automatically to 4.0 or higher by 2 N sodium hydroxide and the agitation was adjusted to up to 1000 rpm dependent on the dissolved oxygen (DO) concentration. A pulse of feed medium (volume corresponding to 30 g C-source per liter initial batch volume) was automatically supplied whenever the C-source was consumed (detected by the increase of DO). Samples for GC-FID analysis of hydrocarbons and fatty acid composition as well as cdw were taken periodically. Samples for the determination of the cdw were centrifuged at 16000 xg, 5 min and dried at 60°C at least 24 h until completely dryness. The cdw was the determined gravimetrically. The physical properties of the LED-strips attached to the bioreactors are summarized in Suppl., Tab. S3.

### 4.6 Lipid extraction, transesterification for GC analysis

For the analysis of lipid content, samples of 1 mL culture volume (or different when indicated) were taken during cultivations and centrifuged at 16000 xg, 5 min. Cell pellets were washed with 1 mL of deionized water followed by a second centrifugation step. Cells were resuspended in 200 µL deionized water and 200 µL glass beads (1:1 mixture of diameters of 0.25-0.5 mm and 0.1 mm) were added to the suspension, as well as 300 µL of n-hexane:2-propanol 3:1 containing internal standard (5 mM tridecanoic acid) for extraction of the triacylglycerols (TAG). Cell lysis was performed in a ball mill (Mixer Mill MM 400) at 30 Hz for 20 min. The lysate was centrifuged at 16000 xg for 1 min and the upper organic phase was transferred to a glass vial. To remove residual water, 50 µL of 2,2-dimethoxypropane were added. Transesterification was performed by the addition of 500 µL 2 % (v/v) methanolic H_2_SO_4_ and incubation at 60 °C and 1400 rpm in an Eppendorf Thermomixer comfort for 2 h. After extraction in 300 µL of n-hexane and optional desiccation over sodium sulfate, the fatty acid methyl ester (FAME) solution was stored at −20 °C until gas chromatography (GC) analysis. For peak assignment, FAME mix from Sigma Aldrich (CRM18918) was used. For quantification, a standard curve of the single FAMEs from Sigma Aldrich Fluka in concentration range of 0.025 - 8 mM were recorded. The samples were analyzed with a Shimadzu Nexis GC 2030, on a Shimadzu SH-Rxi-5MS column (30 m, 0.25 mm, 0.25 µm) and detected by FID. The temperature of inlet and FID were set to 250°C and 310°C, respectively. The linear velocity of hydrogen was set to 50 cm/s. Split to 10. Column oven temperature program: Temp. 90 °C, hold 5 min; Rate 15, final temp. 190 °C; Rate 2.0, final temp. 200 °C, hold 1 min; Rate 0.5, final temp. 202.5 °C, hold 1 min; Rate 20, final temp. 300 °C, hold 5 min. Data were processed using LabSolutions 5.92 and R version 3.4.4 (2018-03-15).

### 4.7 Hydrocarbon analytics of cell extract and supernatant

Analysis was similar to the lipid protocol, including first washing step. Cell lysis was performed, using a Vortexer (10 min, 3000 rpm) from Heathrow Scientific (indicated as non-optimized extraction method in Suppl. Tab. S1/S2). For optimized extraction/lysis, a ball mill (Mixer Mill MM 400) at 30 Hz for 20 min was used. The lysate was centrifuged at 16000 xg for 1 min and the upper organic phase was transferred to a glass vial. For cell extraction, 300 µL n-Hexane containing 5 mM n-dodecane as internal standard, for extraction of whole supernatant (shake flasks experiments) 1.8 mL n-hexane containing 5 mM n-dodecane was used. Detection of hydrocarbons was performed using gas chromatography. GC settings are previously described. The split was set to 50 for samples of total cell extractions from shake flasks, to 5 for 1 ml samples from shake flasks, 24-well and bioreactor measurements and to 10 for samples of the extracted supernatant. The temperature profile was set to an initial temperature of 50 °C, which was held for 2.5 min, followed by a ramp to 250 °C at a rate of 10 °C per min, followed by a ramp to 300°C at a rate of 10 °C per min and a final step at 300°C for 10 min. The analytical GC grade standards undecane, tridecane, pentadecane, heptadecane and the C8-C20 alkane standard solution were purchased at Sigma Aldrich. Quantification of 7-pentadecene, 8-heptadecene and 6,9-heptadecadiene were performed according to [26]. Corresponding peaks were clearly distinguishable from background noise (Suppl., Fig S3 AB). MS spectra, using Shimadzu GCMS QP2010 and column BPX5 (Column has equal properties to SH-Rxi-5MS, but the retention time was slightly shifted, the program is described above) from selected samples were compared to NIST database (GCMSsolution version 4.42, NIST) and confirmed presence of saturated pentadecane (97 % similarity), 8-heptadecene (91 % similarity) and heptadecane (93 % similarity). The retention time difference between monounsaturated 1-pentadecene standard (15.969 min, 98 % similarity) and saturated pentadecane (16.049 min) as well as between 8-heptadecene (18.152 min) and heptadecane (18.433 min) further confirmed the above assumed assignments. Data conversion and plotting was performed with RStudio as described above.

### 4.8 HPLC analysis of extracellular metabolites and media components

Samples were filtered using 10K modified PES Centrifugal filters (VWR). Metabolites of remaining cell free supernatant were analyzed by HPLC. Concentrations of D-glucose, citrate and polyols were determined by Perkin Elmer Series 200, using a RezexTM ROA-Organic Acid H+ (8 %) column (Phenomenex). References were purchased from Sigma Aldrich. The column was eluted with 5 mM sulfuric acid as mobile phase and a flow rate of 0.4 ml min^−1^ at 65 °C. Refractive index values were detected by RI-101 (Shodex). For data evaluation, TotalChrom Workstation/Navigator software (Perkin Elmer, version 6.3.2) was used. Data conversion and plotting was performed with RStudio as described above.

### 4.9 Design and printing of custom labware

CAD for the custom labware was performed using OpenSCAD version 2015.03-1. Except for the designs of the 24-well plates, which was kindly provided by the Möglich lab and is based on their previously published 96-well plates [12]. This design was modified in tinkercad, in order to fit into our microwell holders as well as to accomodate an easier electronic setup by using a FadeCandy (Adafruit Part Number: 1689) driver for the LED-matrix (Adafruit NeoPixel, Adafruit industries, New York, USA). Slicing for 3D printing was performed using Simplify3D version 4.0.1. The labware was printed on a Makergear M2 using PLA as filament.

## List of abbreviations

Å: Ångström
AA: amino acid
AAR (FAR): (fatty) acyl-ACP reductase
ADO: aldehyde-deformylating oxygenase
CAR: carboxylic acid reductase
Cdw: cell dry weight
*Cv/r*: *Chlorella variabilis/reinhardtii*
DO: dissolved oxygen
DOX: α-dioxygenase
FFA: free fatty acids
GMC: glucose-methanol-choline
SCO: single cell oil
FAP: fatty acid photodecarboxylase
FFA: free fatty acids
RT: room temperature
YaliTAR: transformation-associated recombination assisted by *Y. lipolytica*
WT: wild type

## Ethics approval and consent to participate

Not applicable

## Consent for publication

Not applicable

## Availability of data and material

All 3D designs are available in our gitlab repository https://gitlab.com/kabischlab.de/led-labware-cvfap/

## Authors’ Contribution

SB conceived the experimental plan and coordinated the work and analyses. SB and EJM conducted the experimental work. RDL supported the bioreactor cultivations and sample preparations. JK designed and 3D-printed the custom hardware. SB, JK, EJM and RDL prepared the manuscript. RLA provided the strain JMY5749, revised the manuscript and helped with fruitful discussions. All authors read and approved the final manuscript.

## Acknowledgement

We gratefully acknowledge Dr. Mislav Oreb for strain H222 SW1 and Dr. Alexander Rapp for the provision of the broad range photometer. Thanks to Dr. Thomas Hofmeyer for the construction of H222 *Δalk1* strain (S33001), Andreas Möglich for providing the basis for the LED-matrix for microwell plates, Ina Menyes for GC-MS measurements, Felix Melcher for constructing Crispryl-Hyg-URA3 and Belinda Escher for general support.

## Supplementary file

**Tab. S1:**
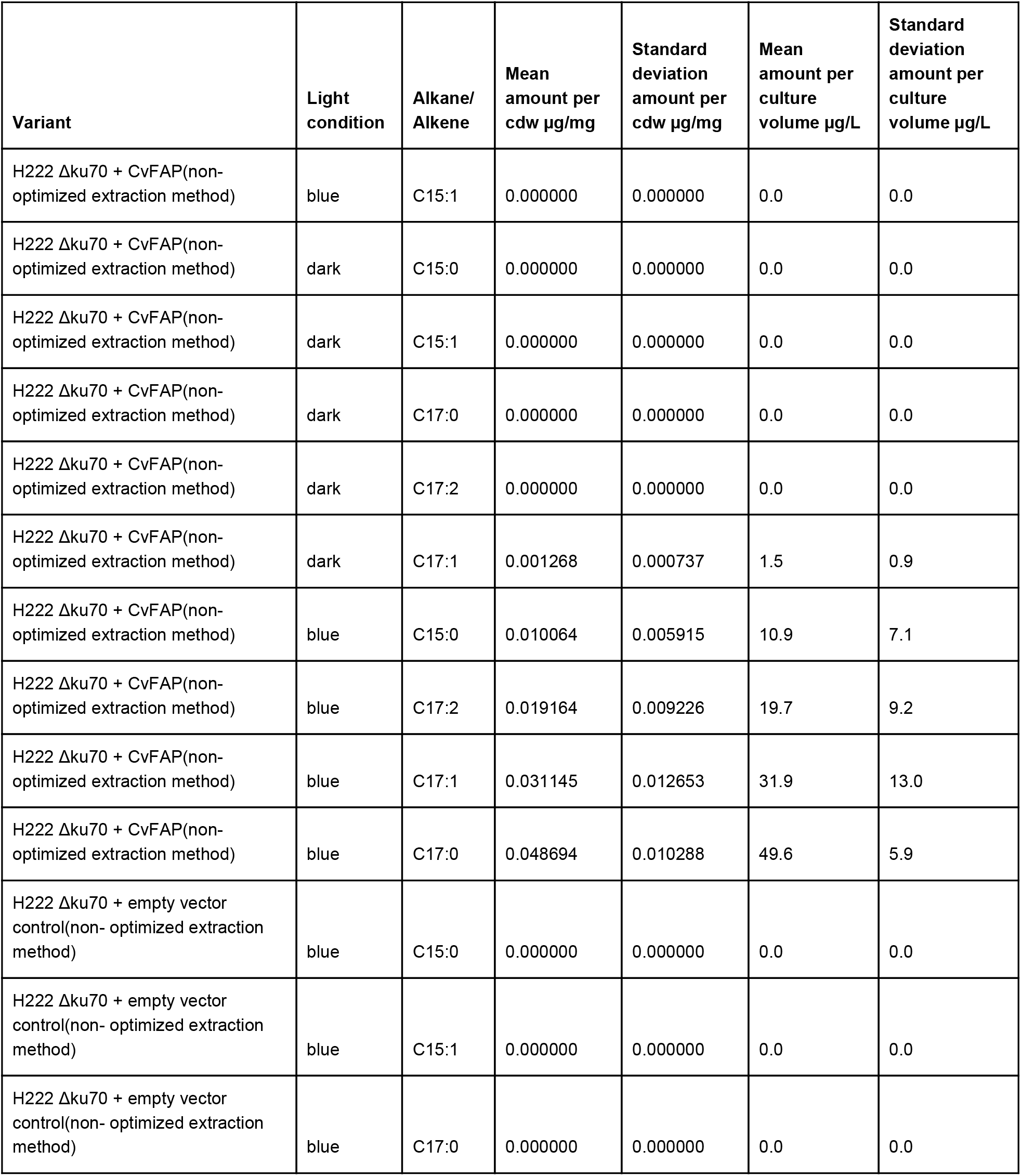
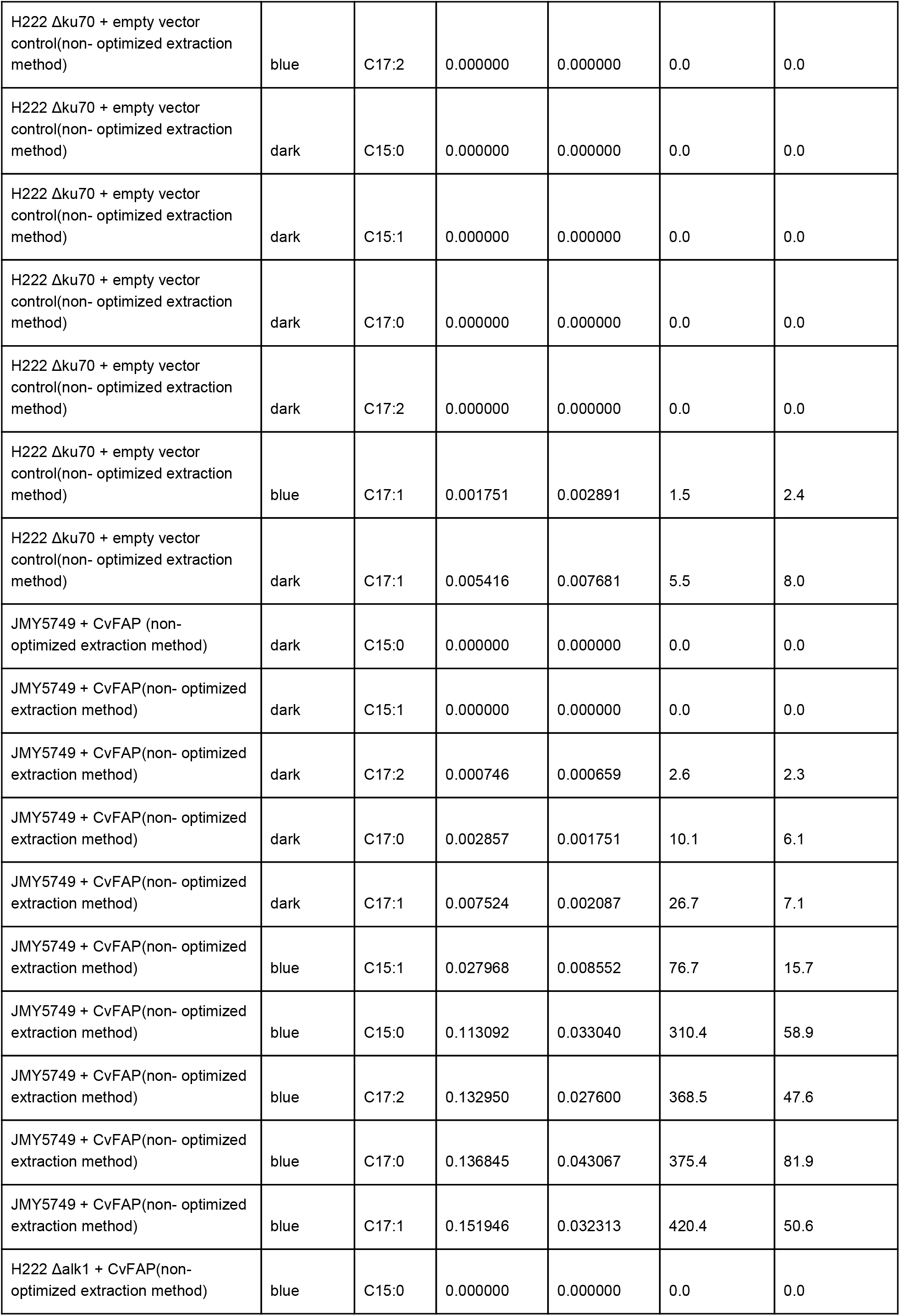
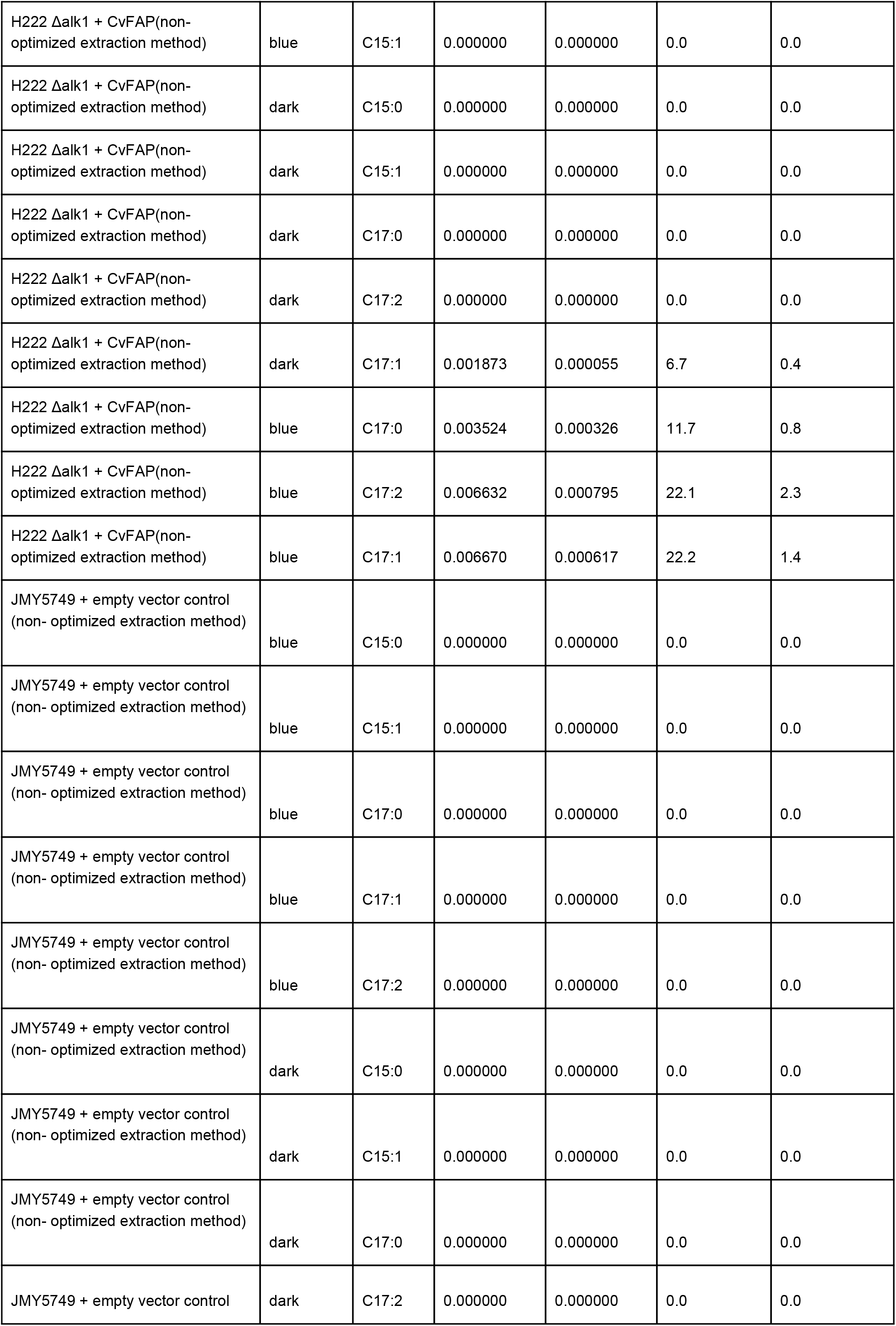
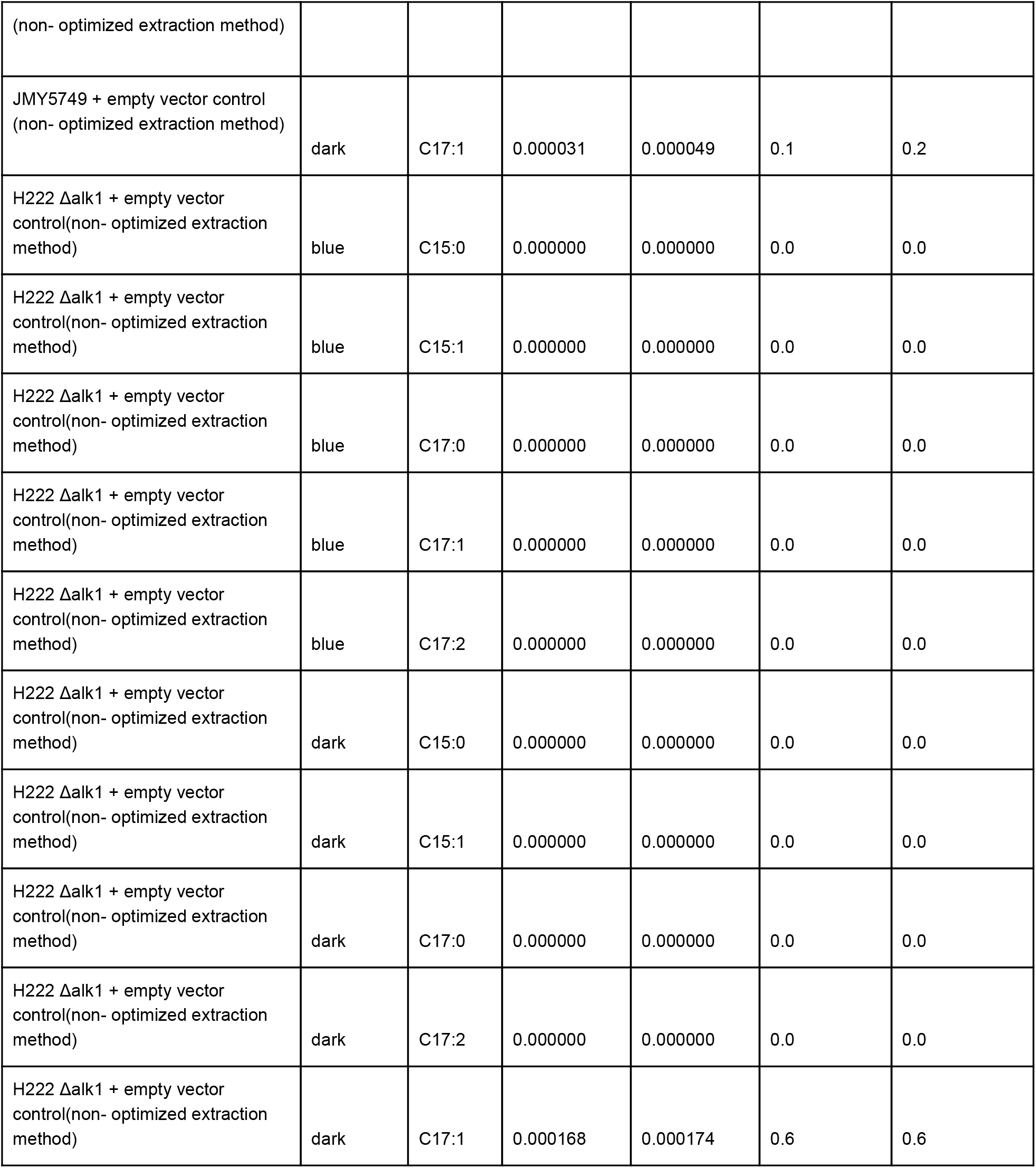

**Tab. S2:**

| **Variant** | **Light** | **Mean cdw**<br>**g/mL** | **Standard**<br>**deviation cdw**<br>**g/mL** | **Mean consumed**<br>**glucose g/L** | **Standard**<br>**deviation**<br>**consumed**<br>**glucose g/L** |
| --- | --- | --- | --- | --- | --- |
| JMY5749 + cvFAP (non- optimized<br>extraction method) | blue | 0.0036 | 0.0005 | 15.6 | 11.1 |
| JMY5749 + cvFAP (non- optimized<br>extraction method) | dark | 0.0040 | 0.0002 | 19.4 | 2.3 |
| H222 Δalk1 + cvFAP(non-optimized extraction method) | blue | 0.0040 | 0.0005 | 16.3 | 5.6 |
| H222 Δalk1 + cvFAP(non-optimized extraction method) | dark | 0.0044 | 0.0008 | 20.9 | 5.9 |
| JMY5749 + empty vector control<br>(non- optimized extraction method) | blue | 0.0049 | 0.0006 | 10.6 | 5.0 |
| JMY5749 + empty vector control<br>(non- optimized extraction method) | dark | 0.0047 | 0.0007 | 13.3 | 3.8 |
| H222 Δalk1 + empty vector<br>control(non- optimized extraction<br>method) | blue | 0.0034 | 0.0004 | 23.8 | 2.4 |
| H222 Δalk1 + empty vector<br>control(non- optimized extraction<br>method) | dark | 0.0069 | 0.0016 | 11.4 | 11.8 |

**Tab. S3:** Correlation light intensity / power supply values. Single LED measurements to determine technical conditions for bioprocess cultivation. The applied voltage was limited to a maximum of 12 V. The LED-strip was connected to the power-supply Consort EV231 (purchased from Sigma Aldrich). Photosynthetically active radiation (PAR) was measured using a Li-190SA quantum sensor (Li-COR) coupled with a Li-1000 (Datalogger).

| Intensity<br>[$\mu\text{mol photons s}^{-1} \text{m}^{-2}$] | Amperage<br>[mA] | Voltage<br>[V] | Measuring Error<br>± |
| --- | --- | --- | --- |
| 68 | 100 | 2 | 8 |
| 130 | 200 | 2 | 9 |
| 192 | 300 | 3 | 9 |
| 250 | 400 | 2 | 9 |
| 310 | 500 | 2 | 9 |
| 357 | 600 | 1 | 10 |
| 415 | 700 | 2 | 10 |
| 455 | 800 | 5 | 11 |
| 533 | 900 | 5 | 11 |
| 545 | 1000 | 5 | 12 |

**Tab. S4:**
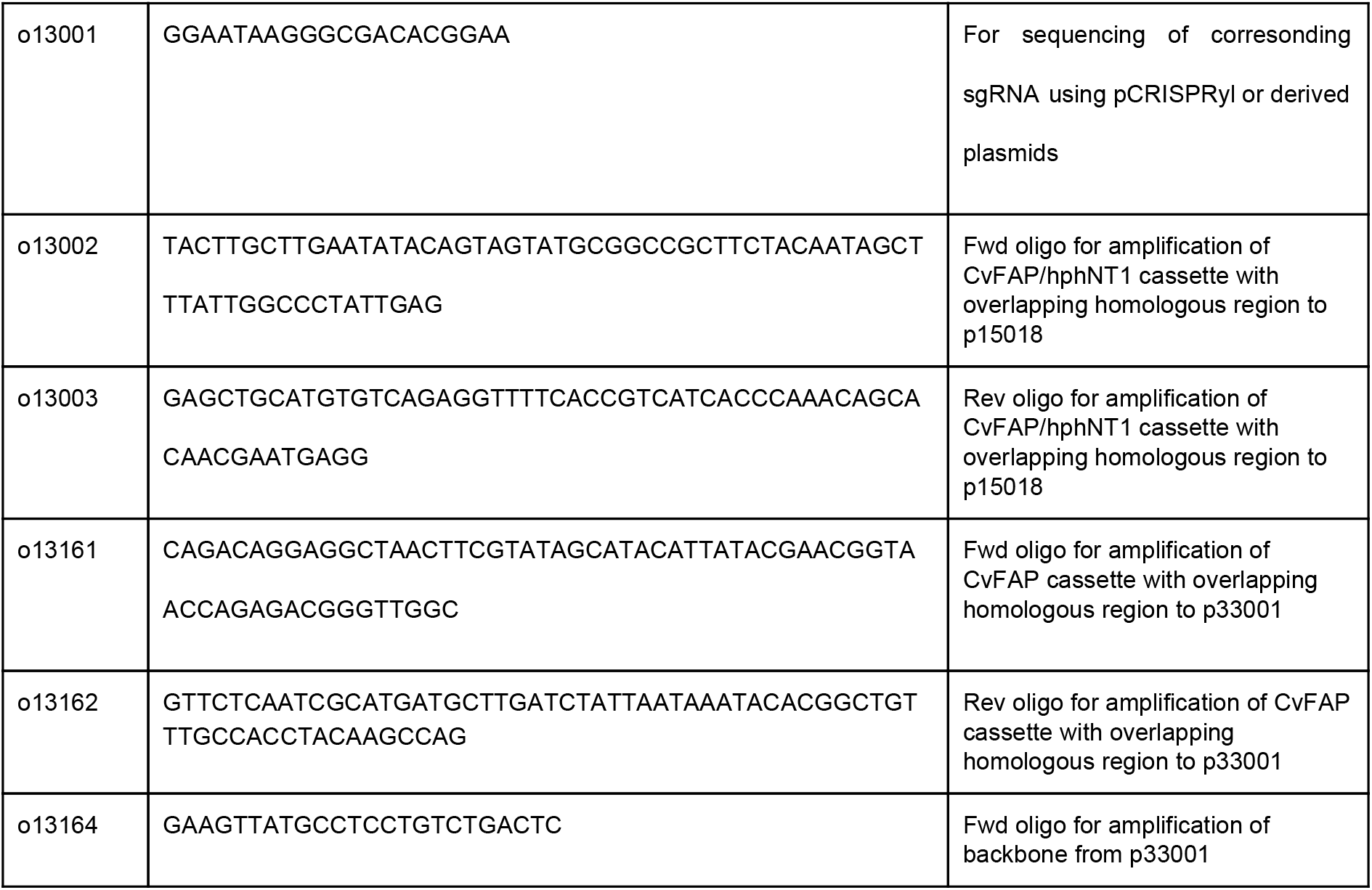
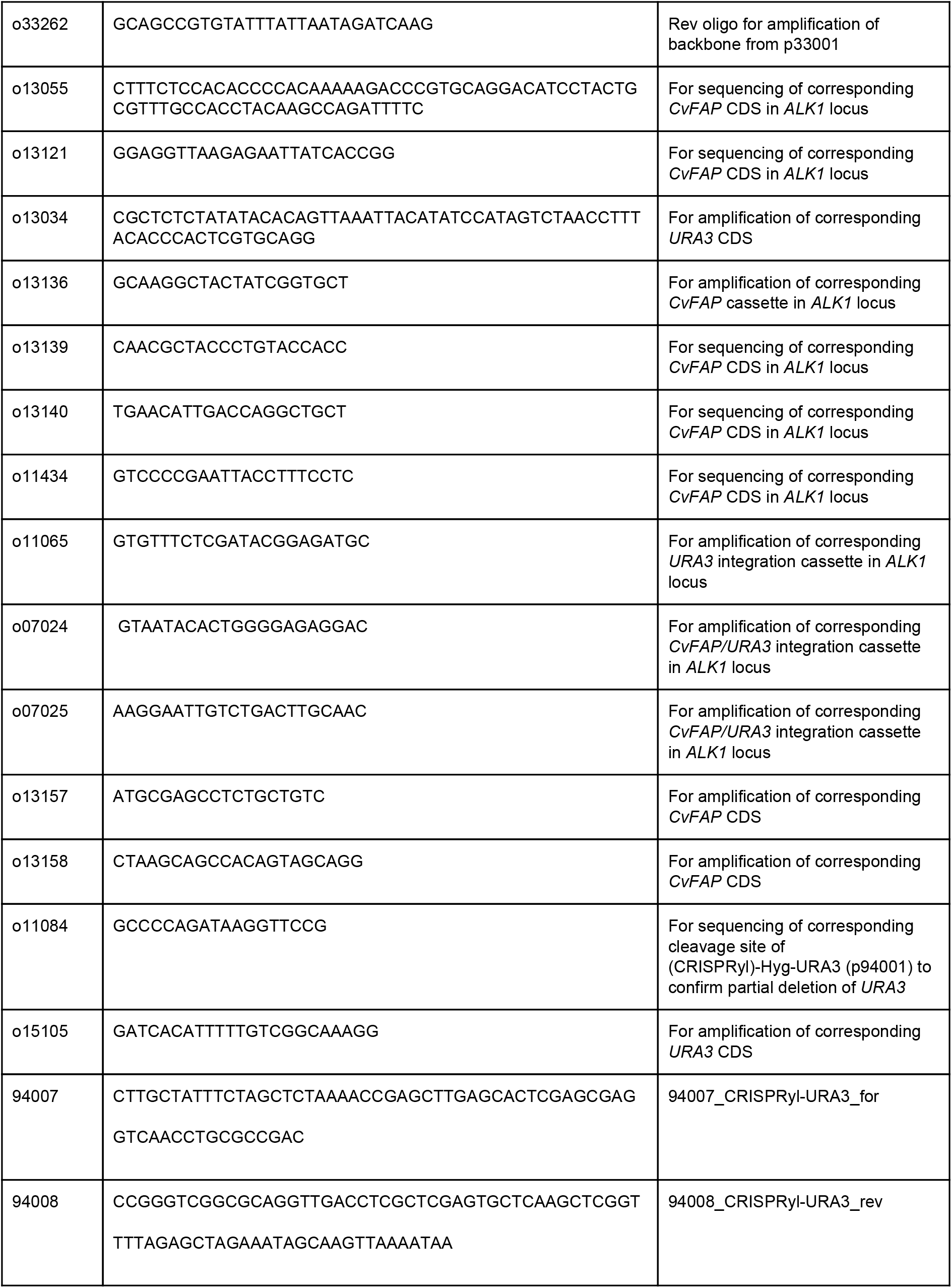
List of oligonucleotides.

**Tab. S5:**
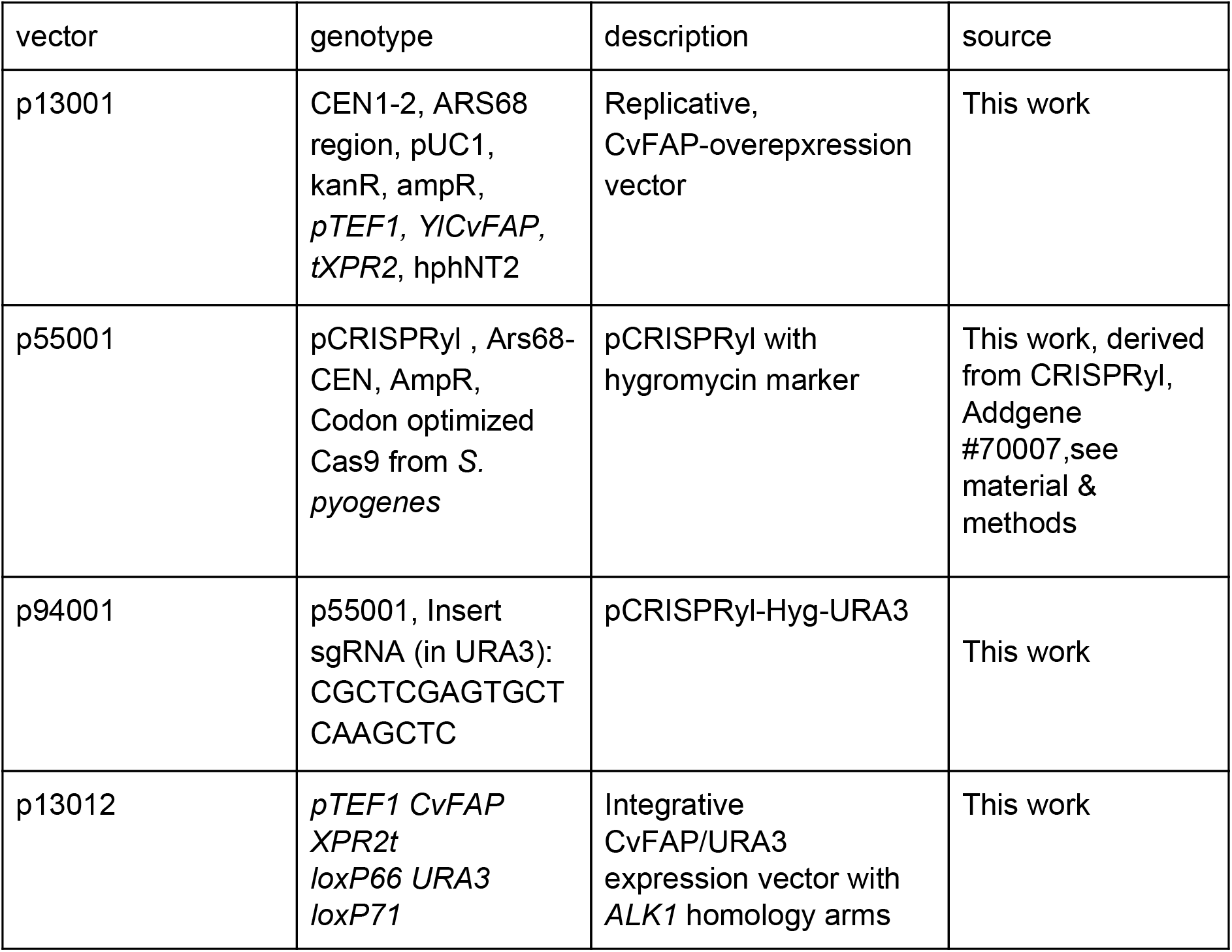
Constructed vectors.

**Fig. S1:**
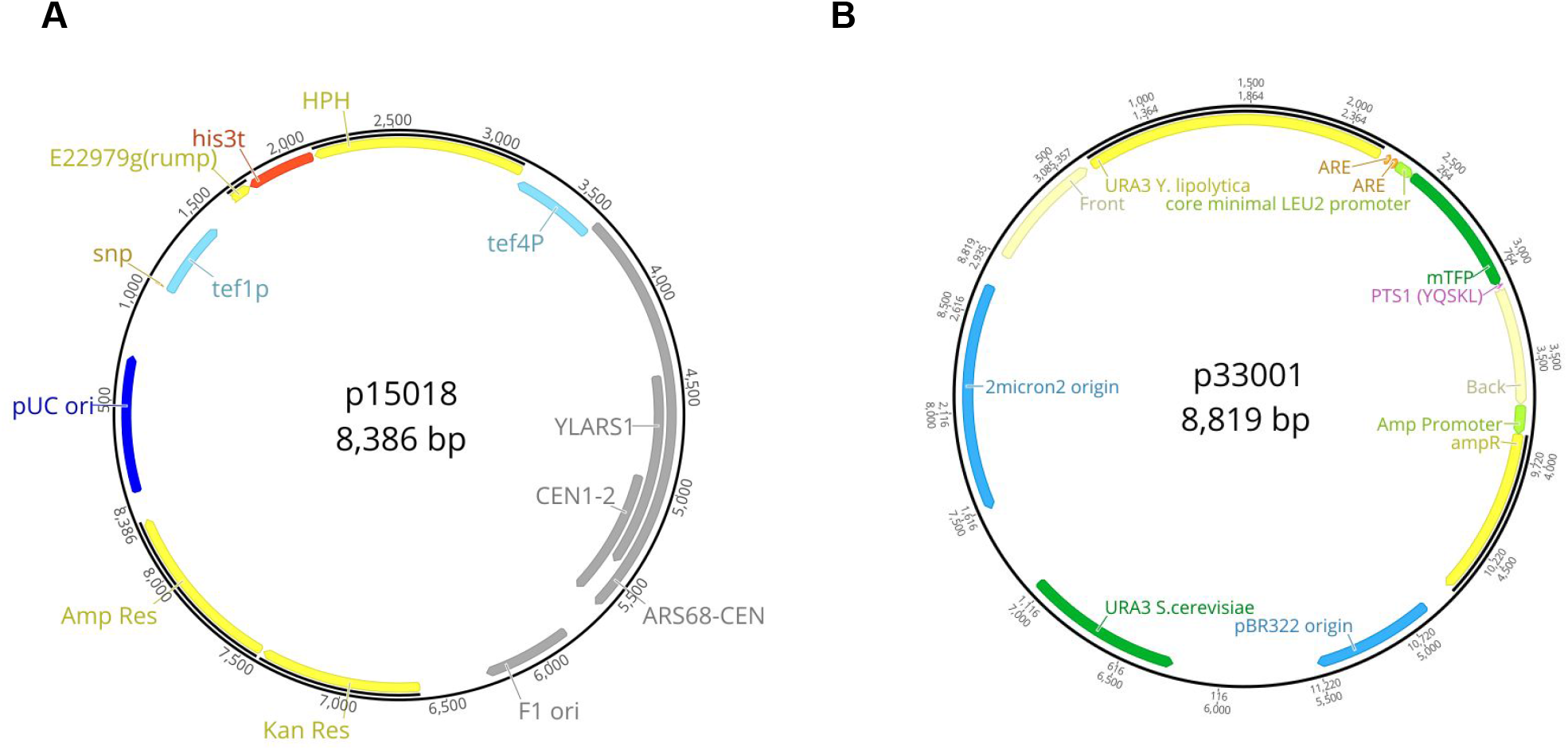
Vector maps were created by using Geneious v.10.2.3. **A:** The replicative, centromeric *Y. lipolytica, E. coli* shuttle-plasmid p15018 consists of CEN1-2, ARS68 region, pUC1 ori, kanamycin, ampicillin resistance cassette, as well as native *pTEF1* promoter and *tXPR2* terminator for gene expression and hphNT2 cassette (HPH) encoding for hygromycin resistance. **B:** The replicative, centromeric *S.cerevisiae, E. coli* shuttle-plasmid p33001 serves as integration vector in *Y.lipolytica*. It consists of 2µ region, *Sc*URA3 marker gene, pBR322 origin and ampR marker. Front and Back region are homologous to promoter and terminator region of *YlALK1*.

**Fig. S2:**
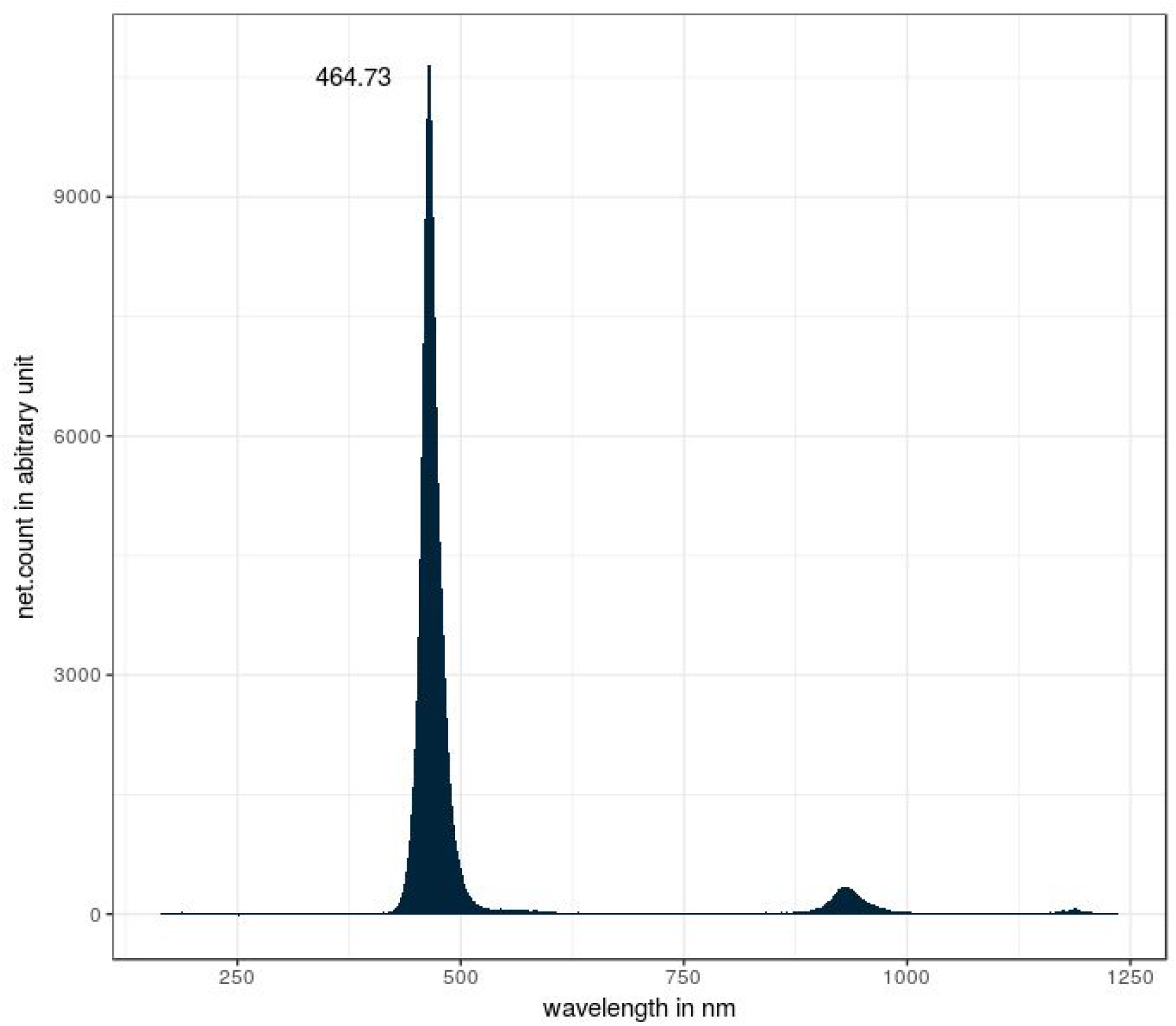
Light emission of advertised distinct 465-470 nm LED-light strip by wavelengths. The defined spectrum was verified by broad range spectrometer LR1-T from ASEQ instruments.

**Fig. S3.**
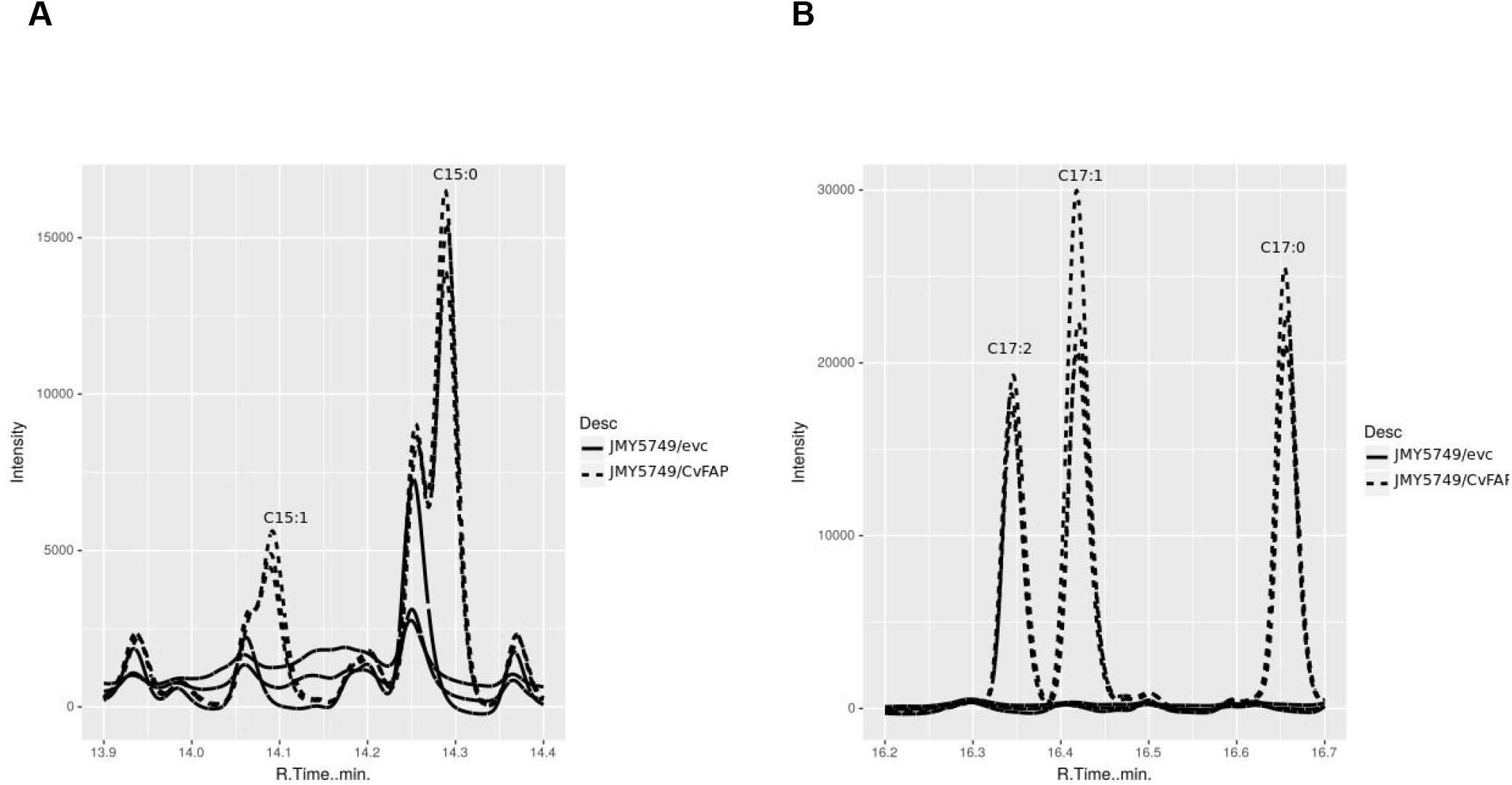
**A:** Assigned peaks of pentadecane (C15:0), 7-pentadecene (C15:1), **B:** heptadecane (C17:0) 8-heptadecene (C17:1) and 6,9-heptadecadiene (C17:2) are shown for analysis of cell extraction of alkane producing strain JMY5749/CvFAP in comparison to the empty vector control.

**Fig. S4:**
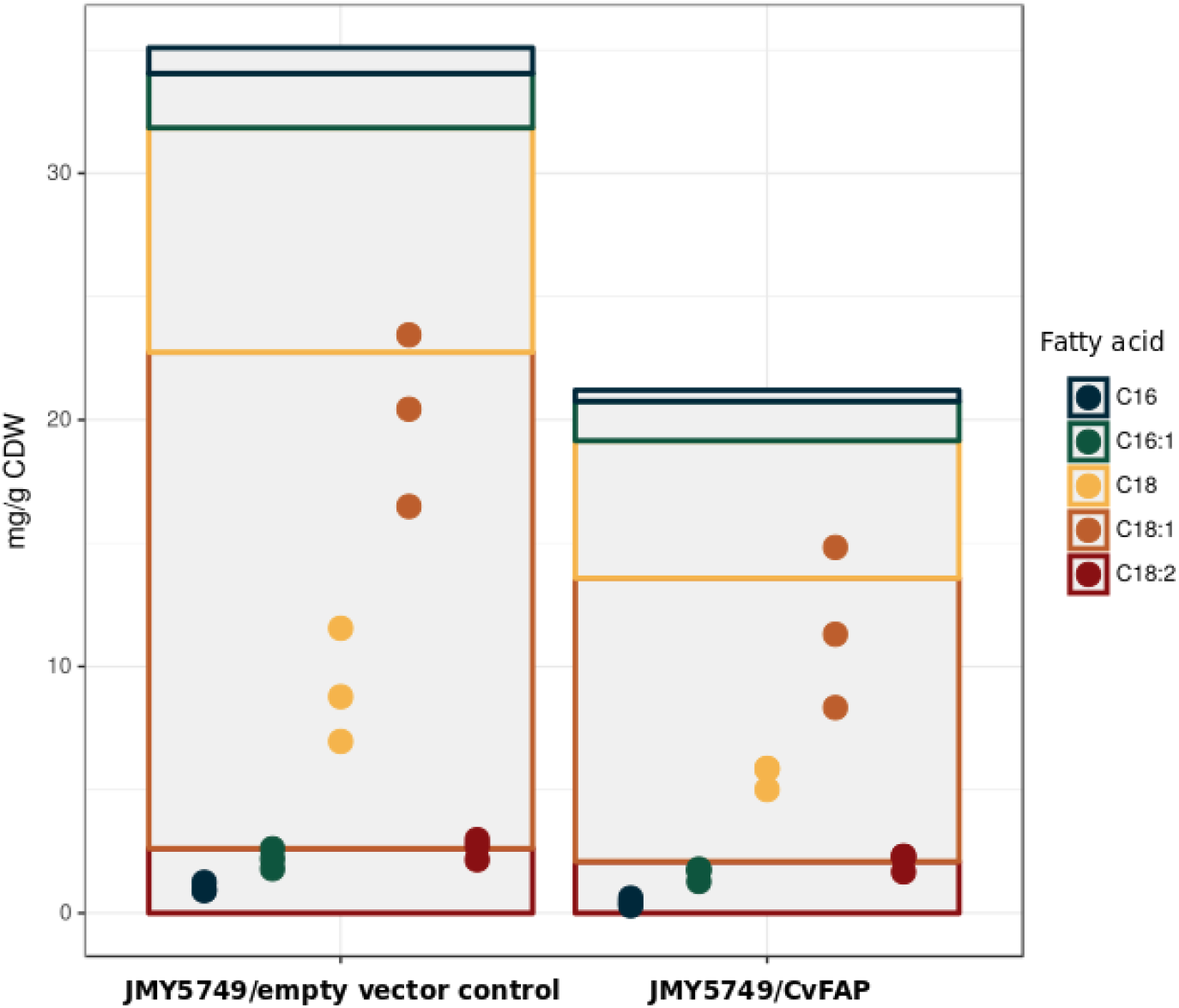
Intracellular fatty acid concentrations of *Yarrowia* JMY5749/CvFAP constructs and empty vector control. Cultivation conditions are listed in publication Fig.2 and above. Description of lipid analytics is mentioned below. The concentration of each fatty acid is indicated by dots (in triplicates), the sum (medium) of fatty acids and the fatty acid composition is represented by a bar plot. Total fatty acid concentration was significant lower (Students two Sample t-test: t=3.2778, df=4, p-value=0.03056) for JMY5749/CvFAP construct (21.20272+/-4.396019) in comparison to empty vector control (35.08924+/-5.875347).

**Fig. S5:**
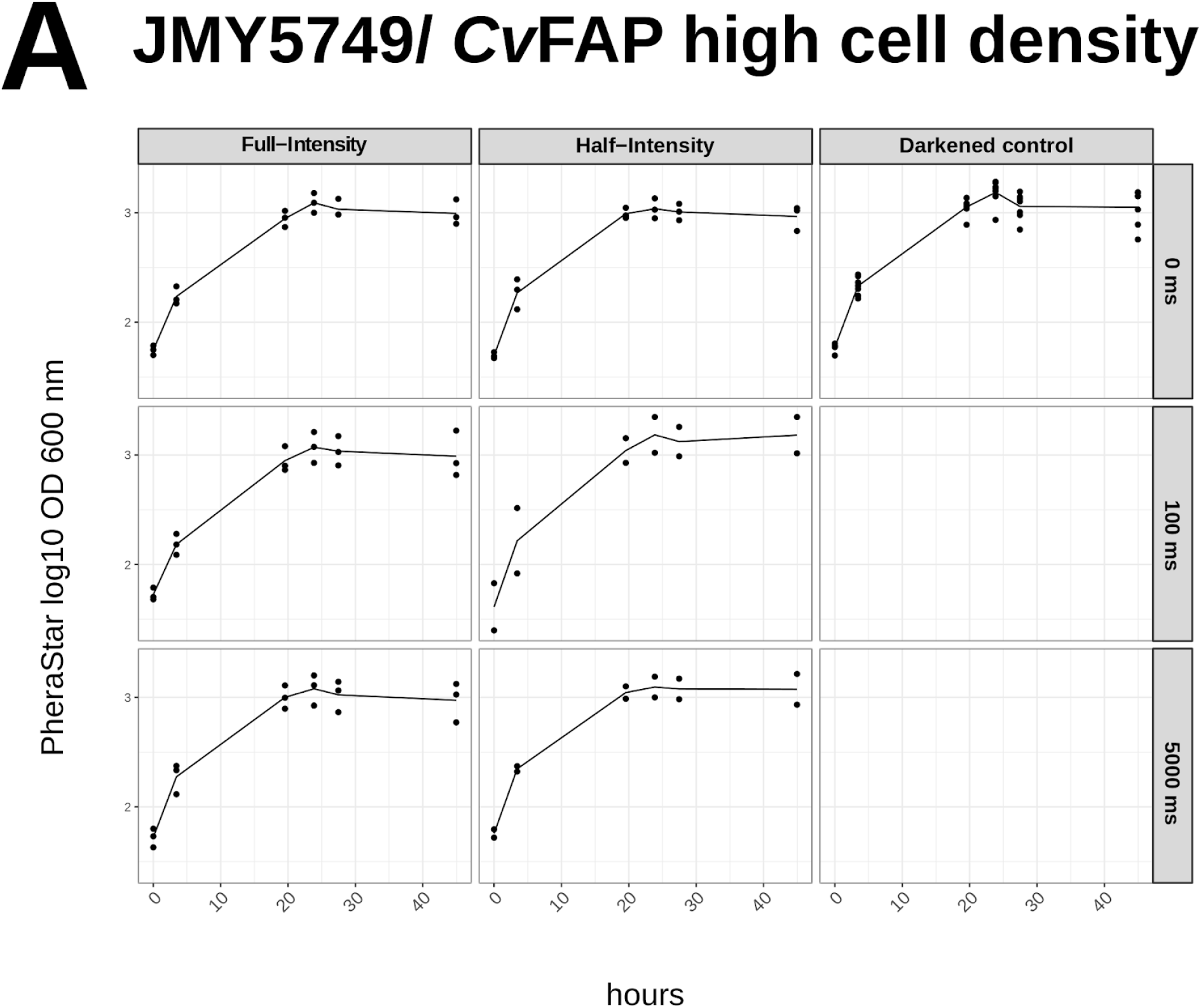
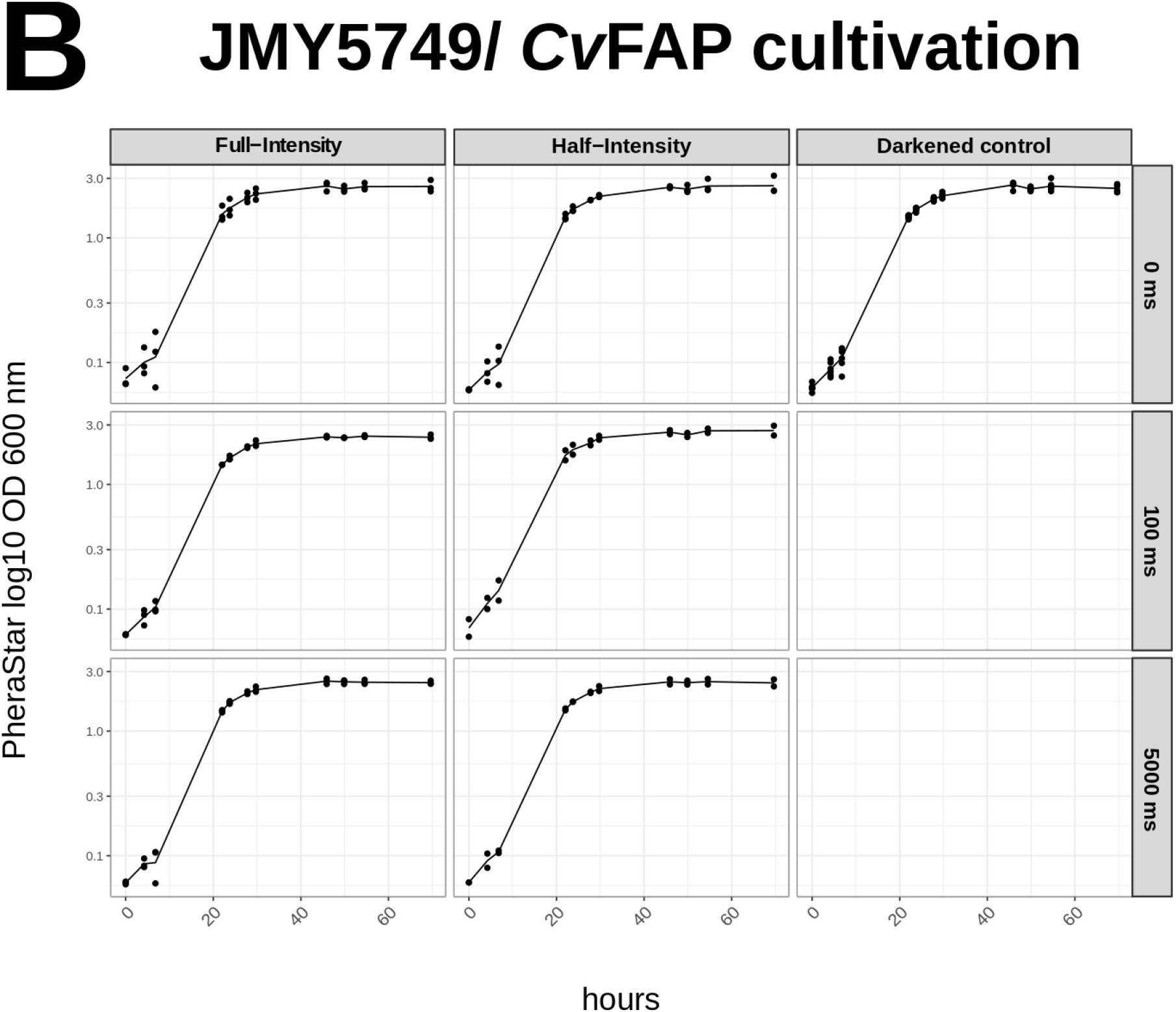
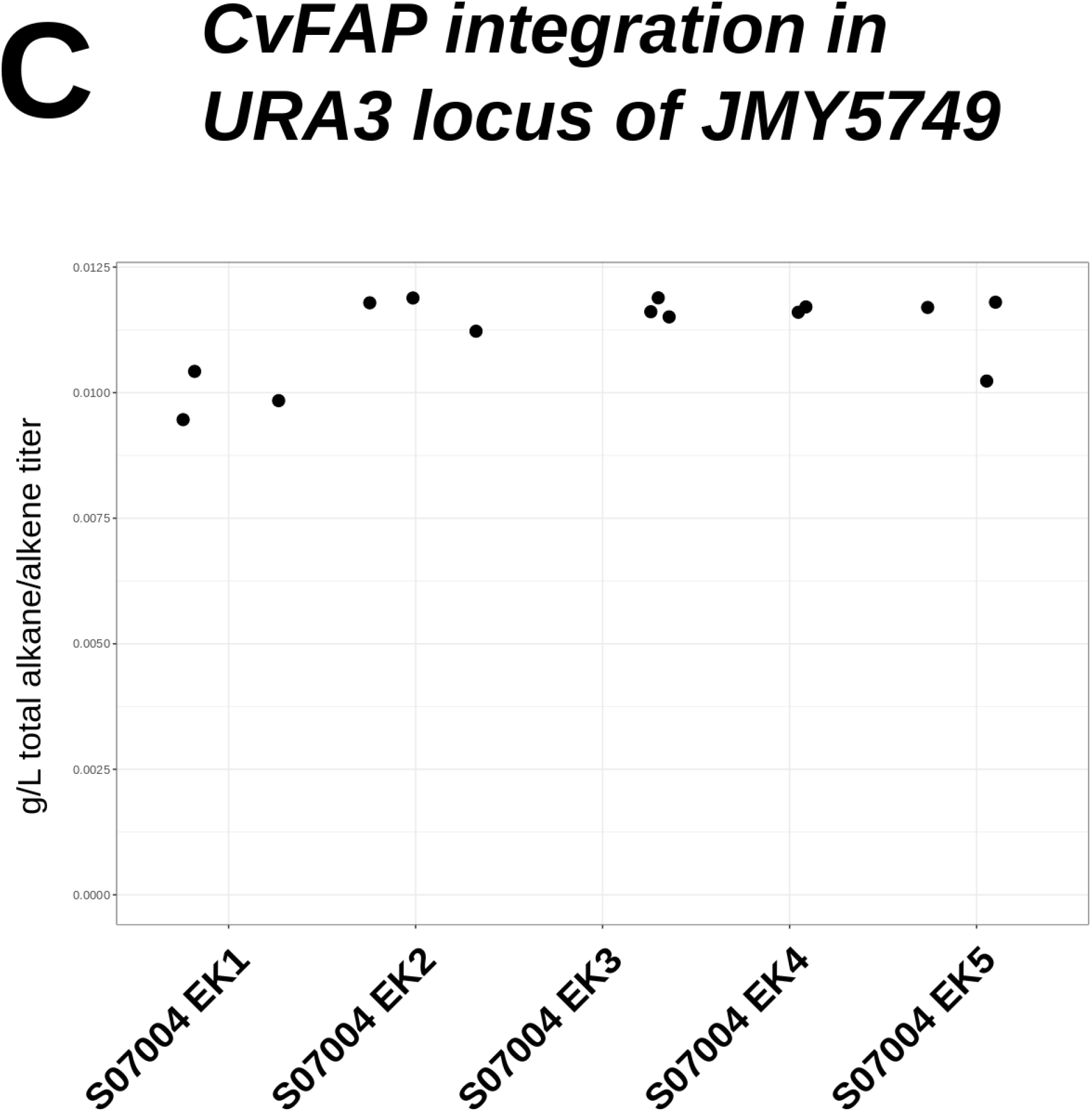
A: Time-resolved logarithmized optical density measurements of light biotransformations, shown in Figure 3 A. B: In opposite to high cell density biotransformation (A), initial OD_600_ were set to 0.1 to determine the effect of different light intensities on cell growth. Light regimes were tested in triplicates, except for Half-Intensity, pulse 100 and 5000 ms. C: Endpoint measurements of total hydrocarbon formation for cultivations (initial OD_600_ 0.1) of different S07004 clones, harbouring the genomic integration of *Cv*FAP cassette. Strains were cultivated in triplicates, expect for S07004 EK4. Corresponding alkane/alkene composition for endpoint measurements are shown in Figure 3 C.

**Fig. S6:**
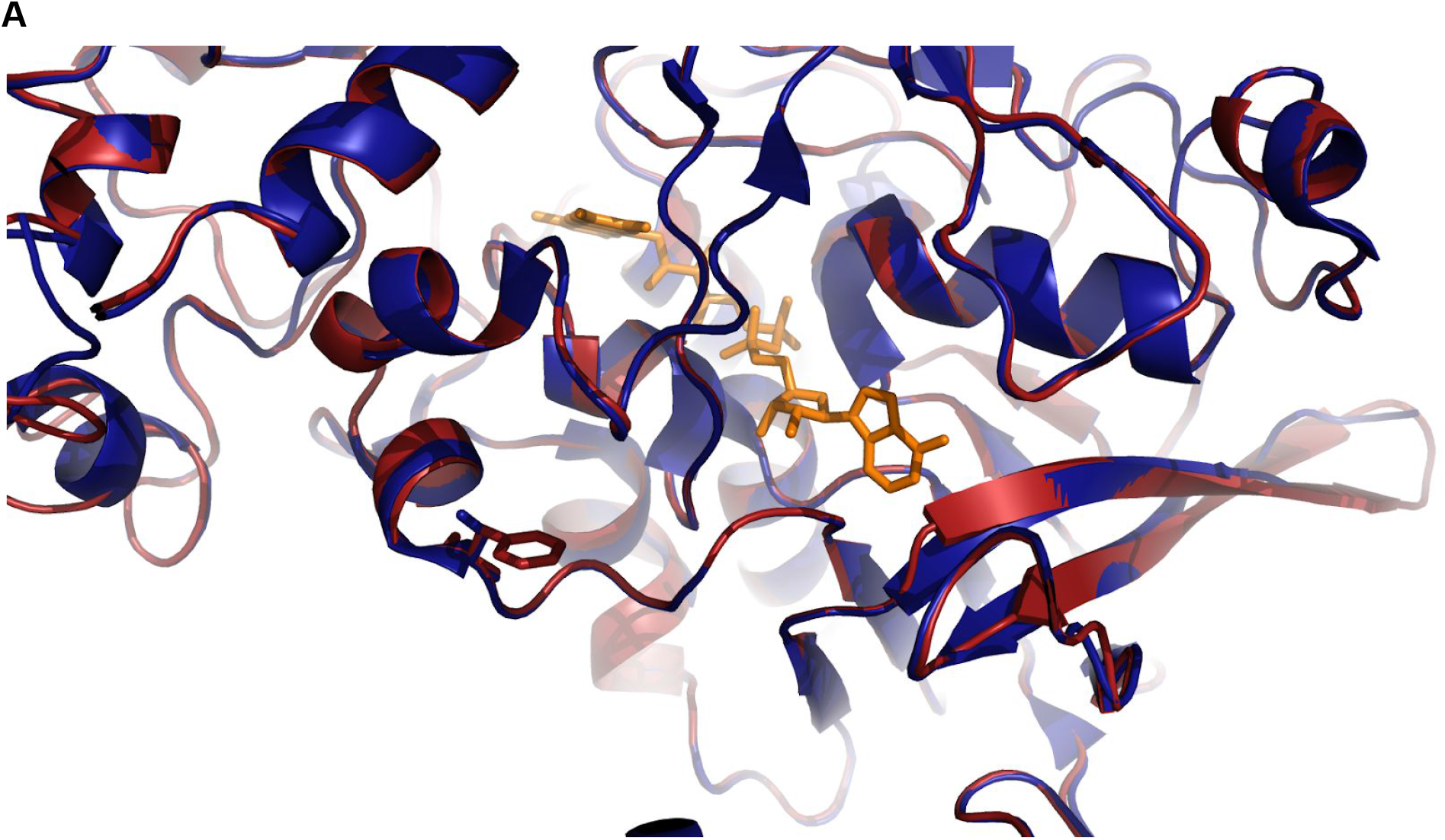
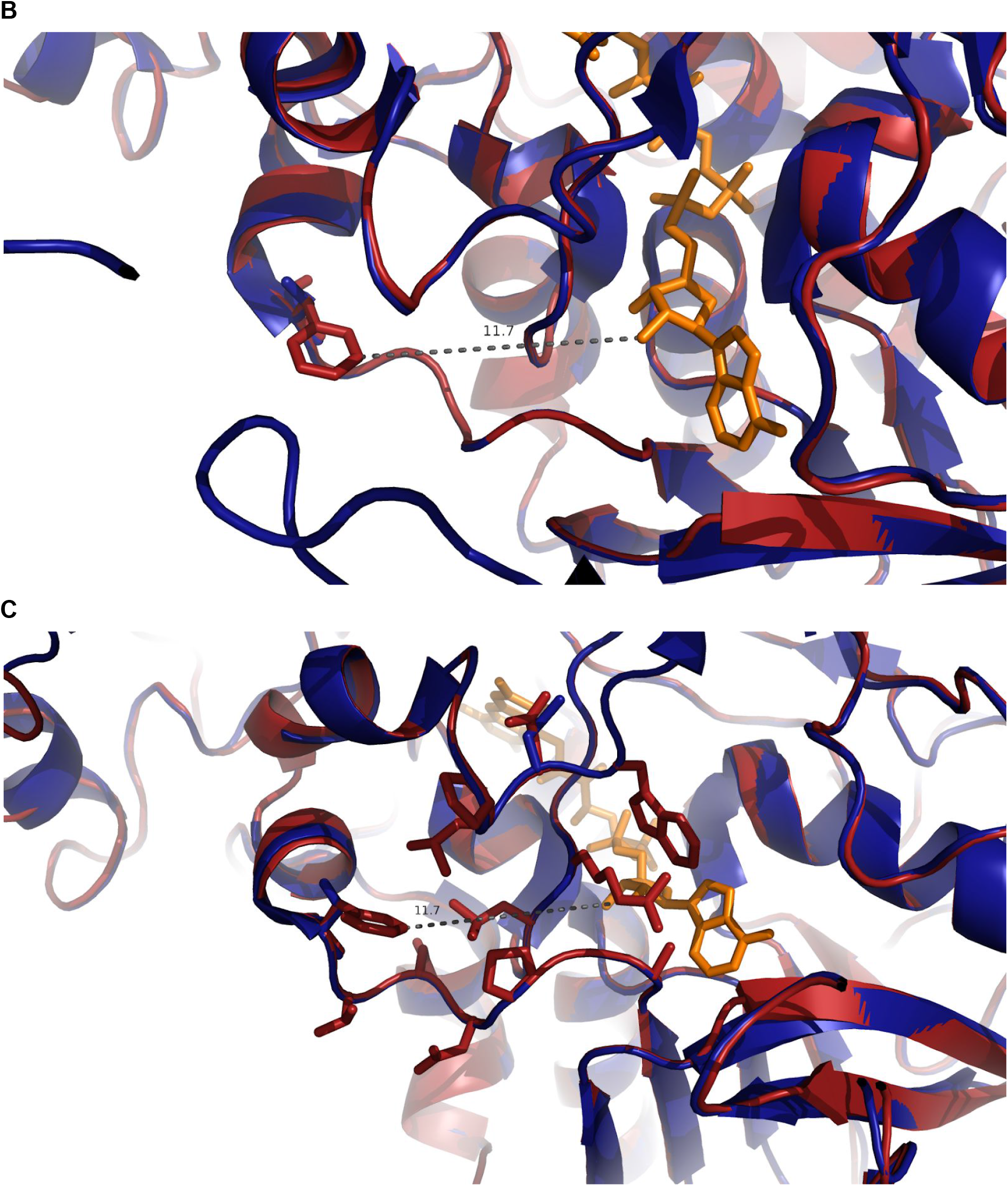
Rendering of CvFAP WT (blue) and S61F mutant (red) using PyMOL 1.7.x. Homology model of the mutant was created by using phyre2 algorithm [1] and pdb structure of the wild type (5NCC), deposited by [2]. A: Overview of cofactor binding side; B: Minimum distance in Ångström, between AA 61 (121 for WT CvFAP) phenylalanine ring and flavin adenine dinucleotide; C: Amino acid residues in close proximity to residue 121 and FAD.

**Fig S7:**
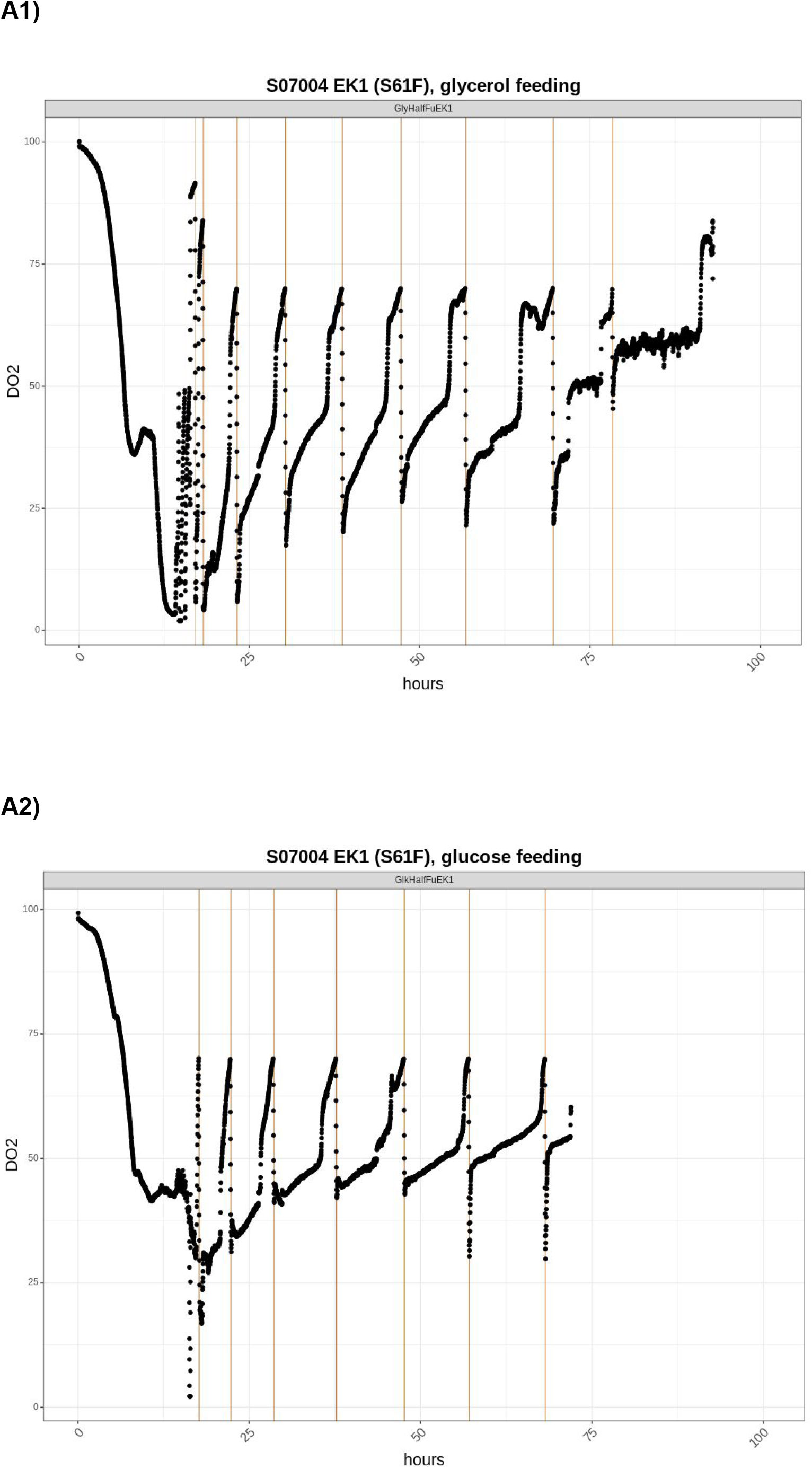
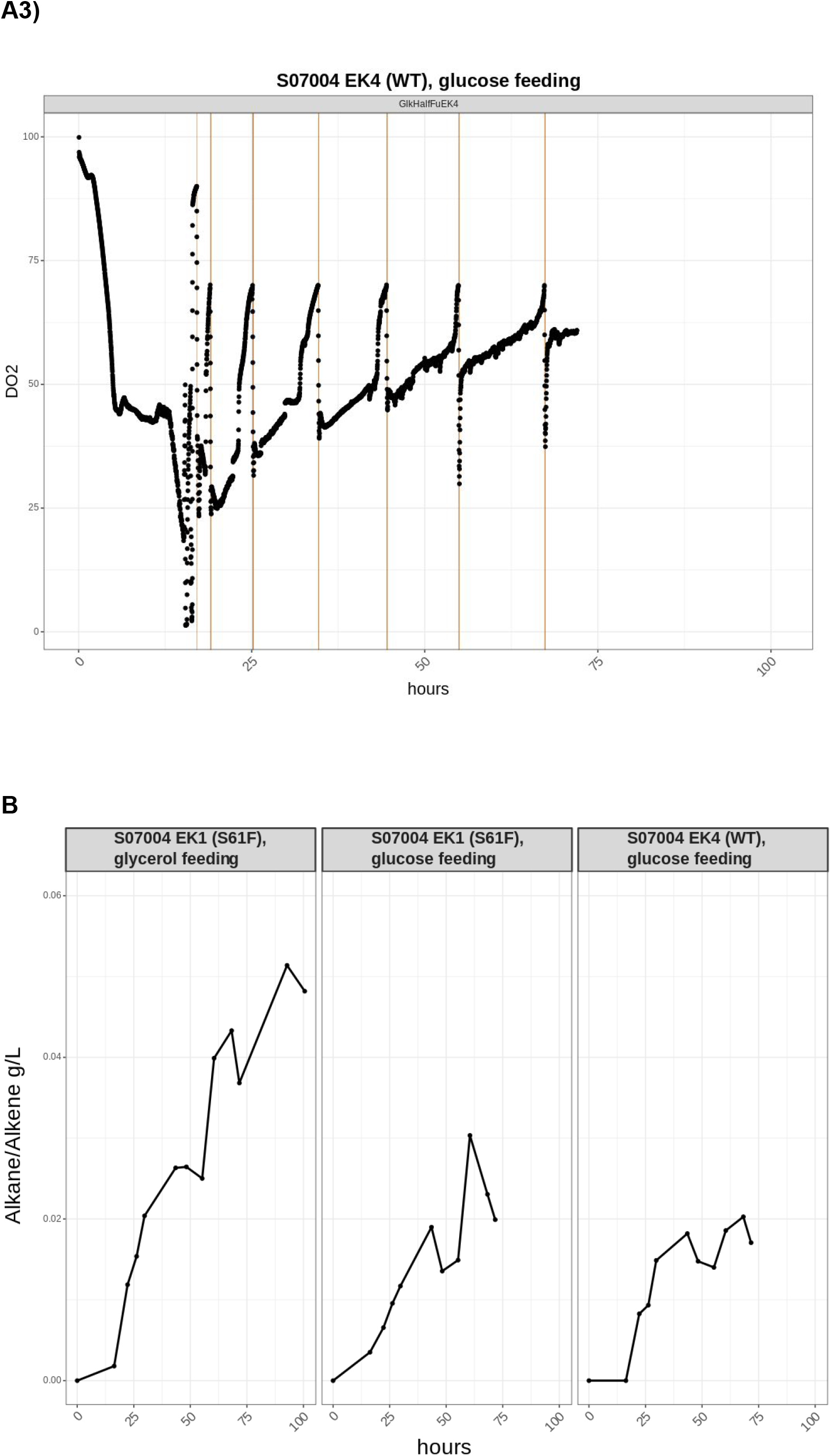
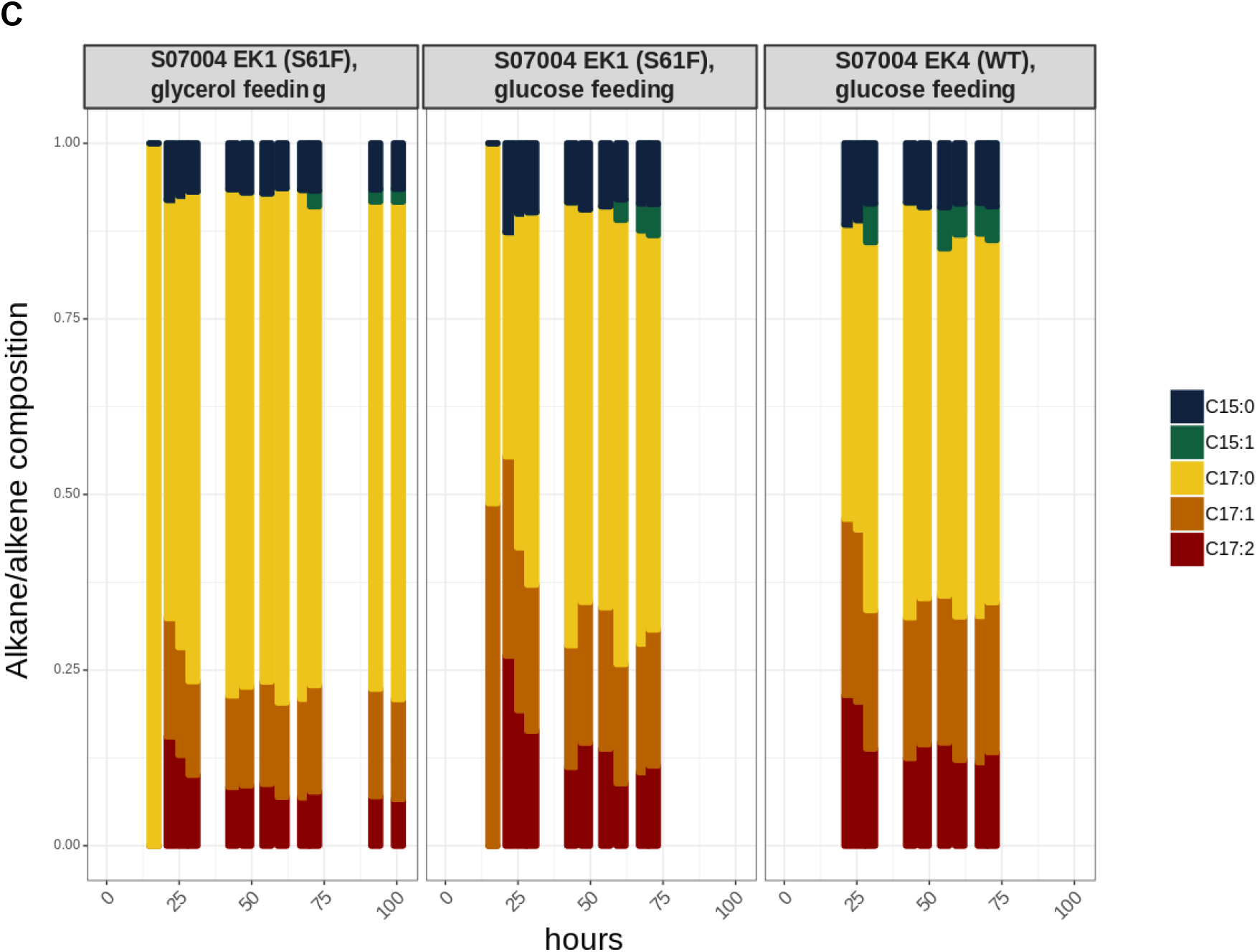
Light settings: Half-Intensity after inoculation, Full-Intensity after 22.25h A1)

**Fig S8:**
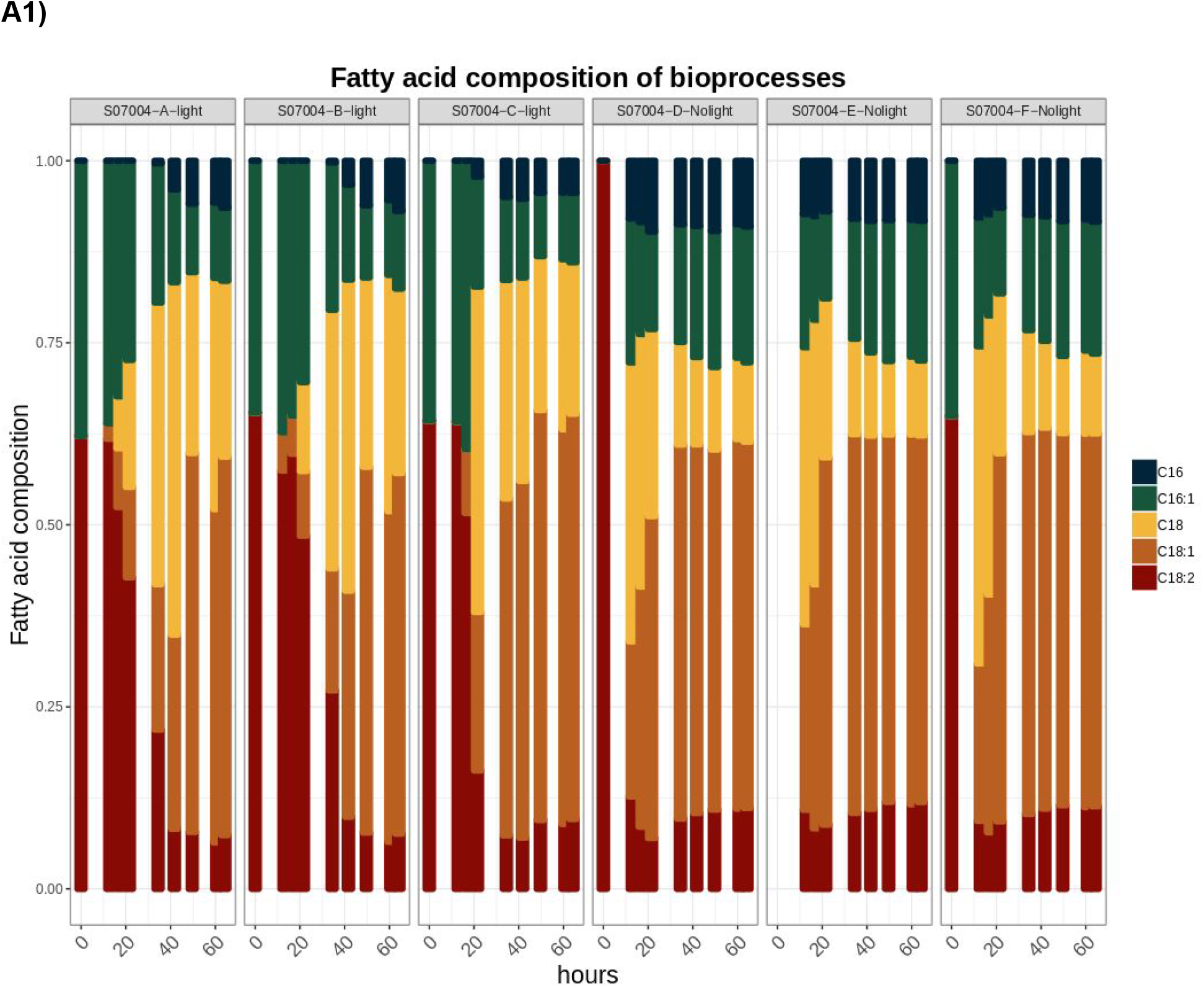
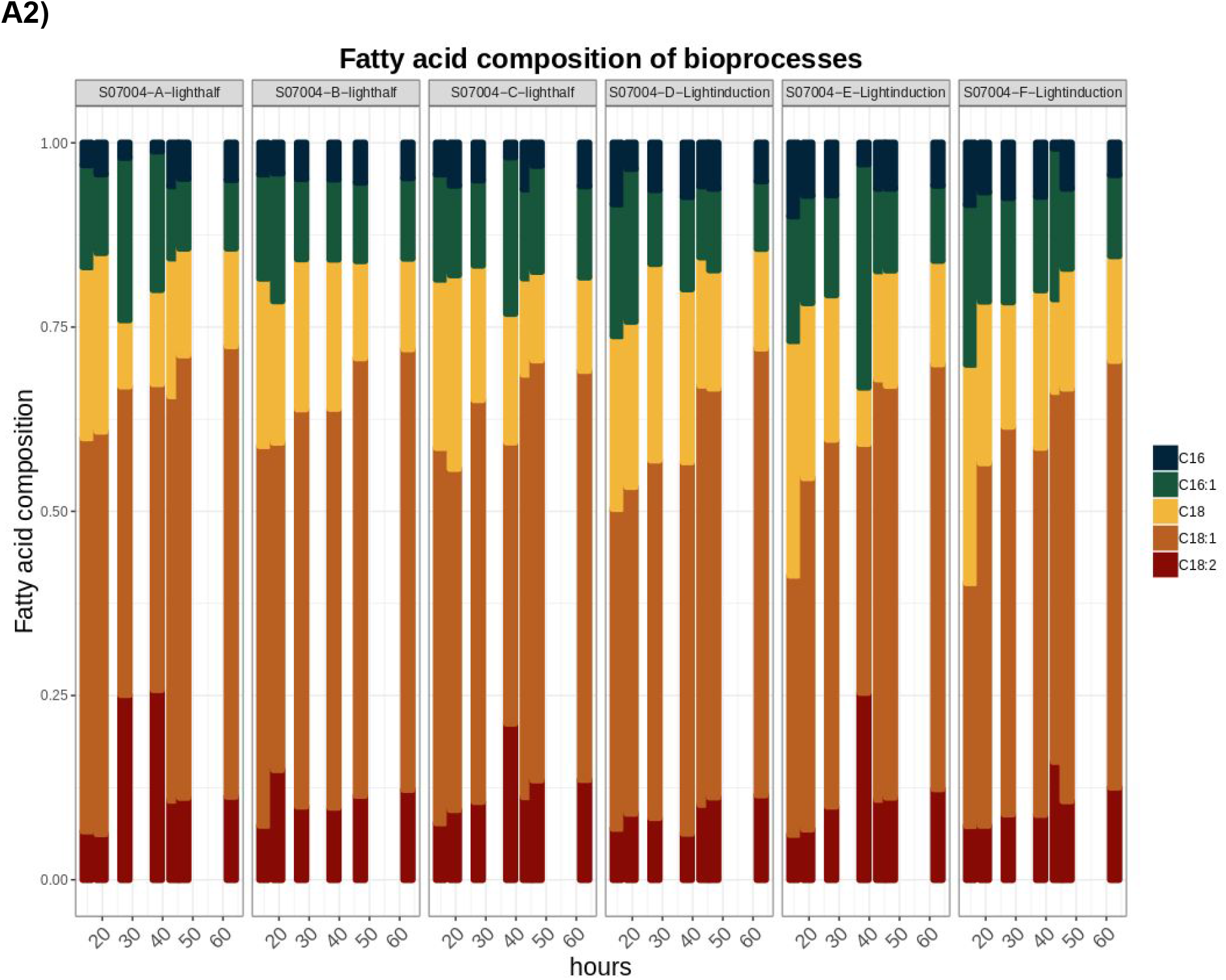
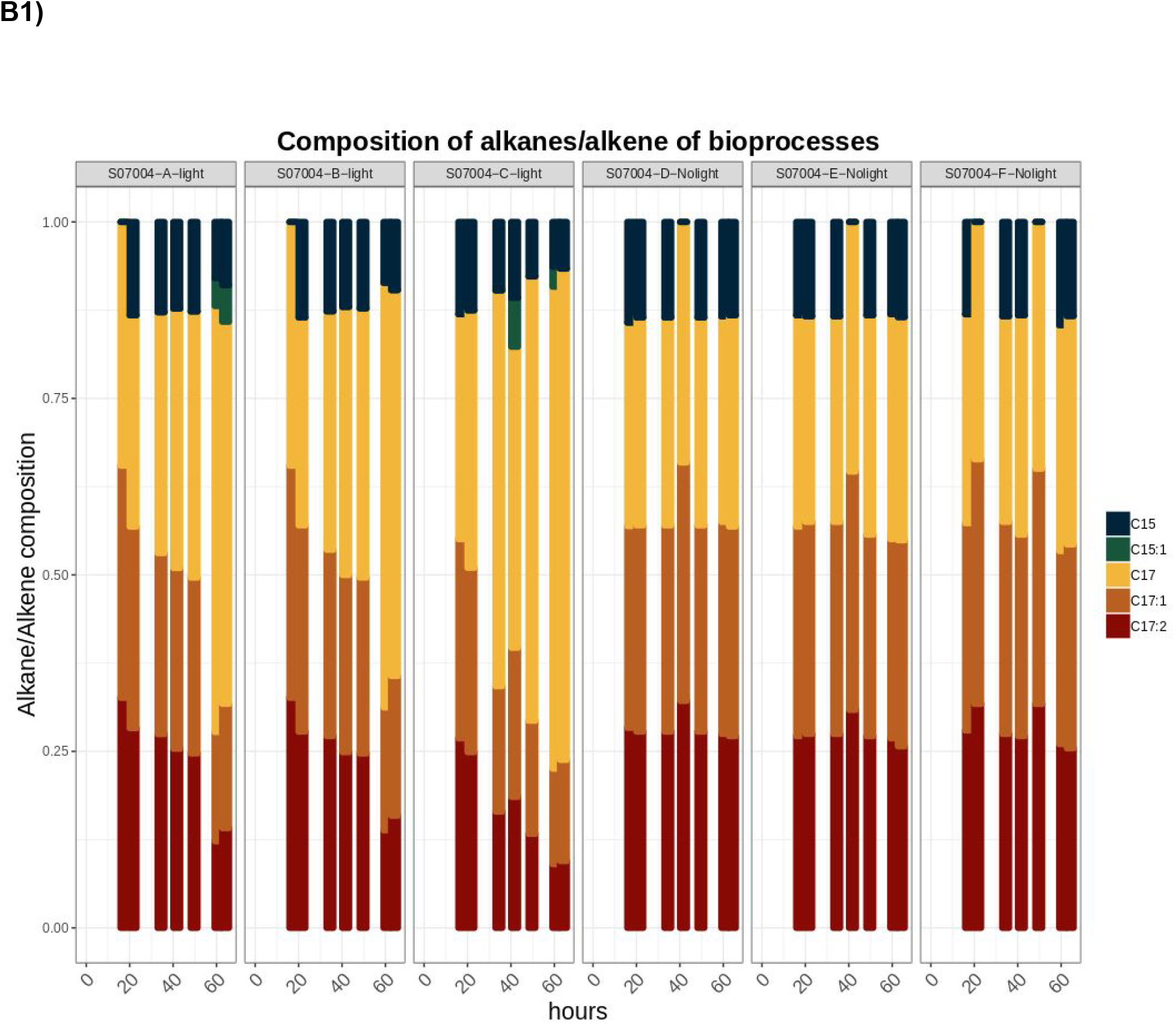
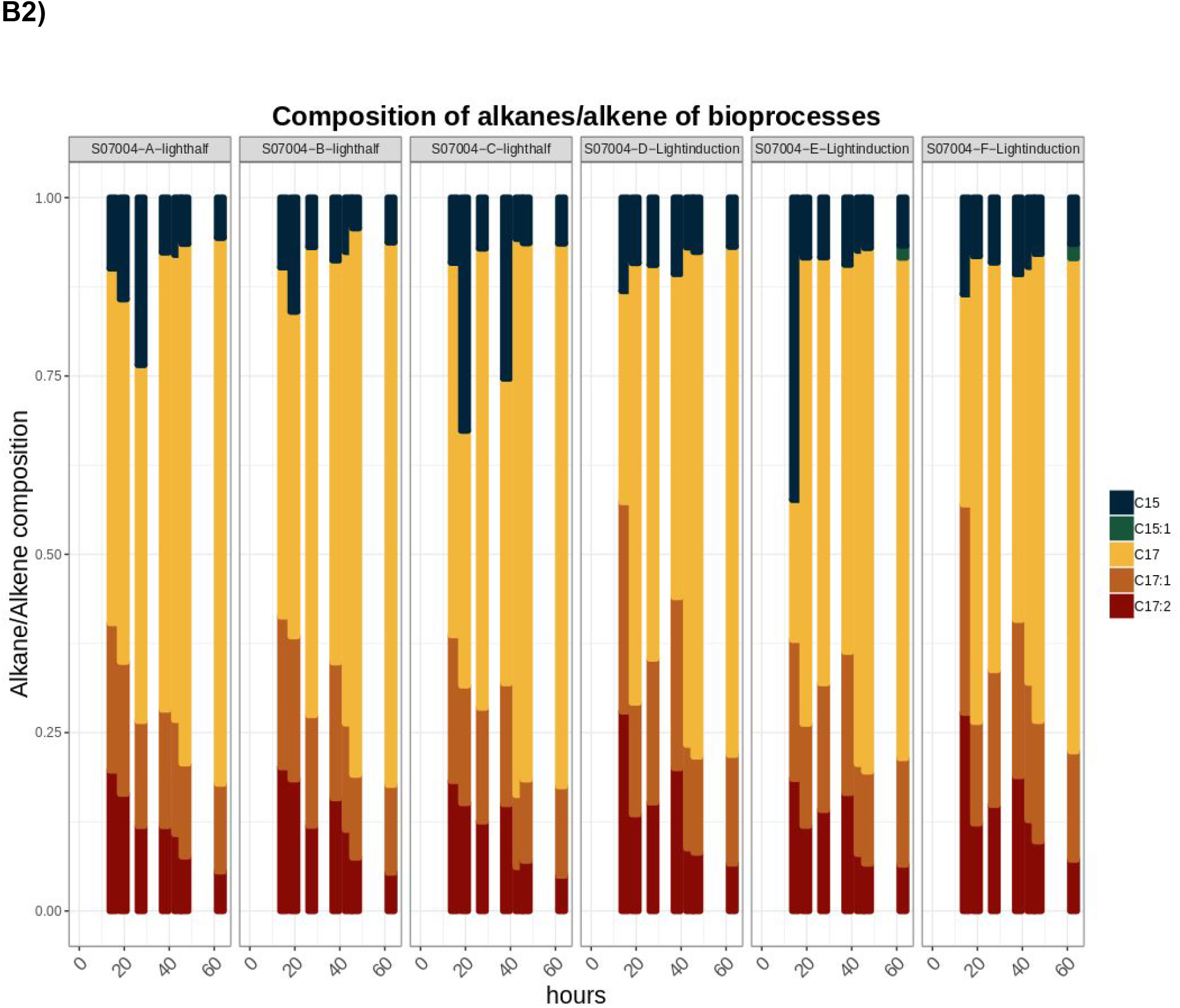
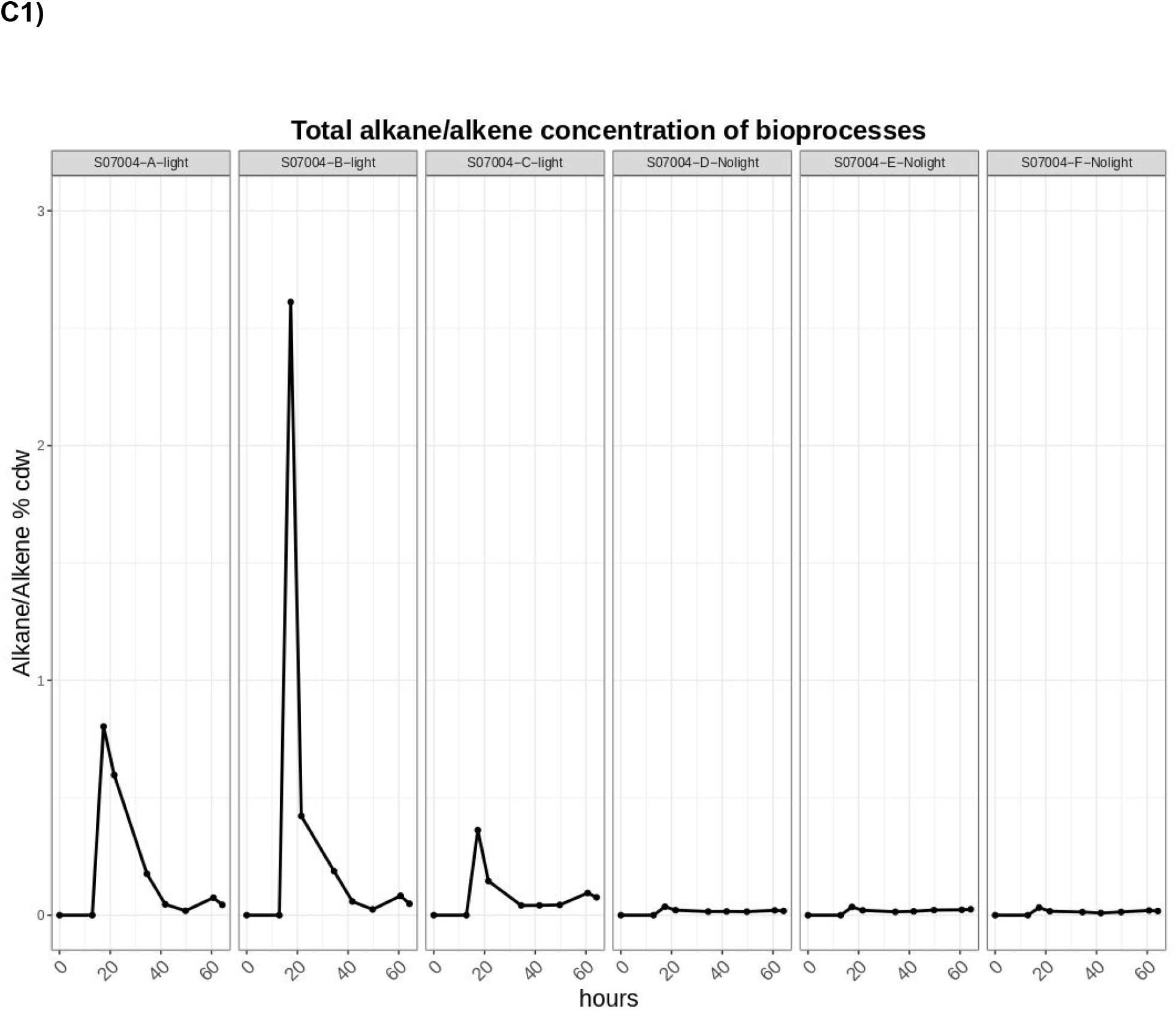
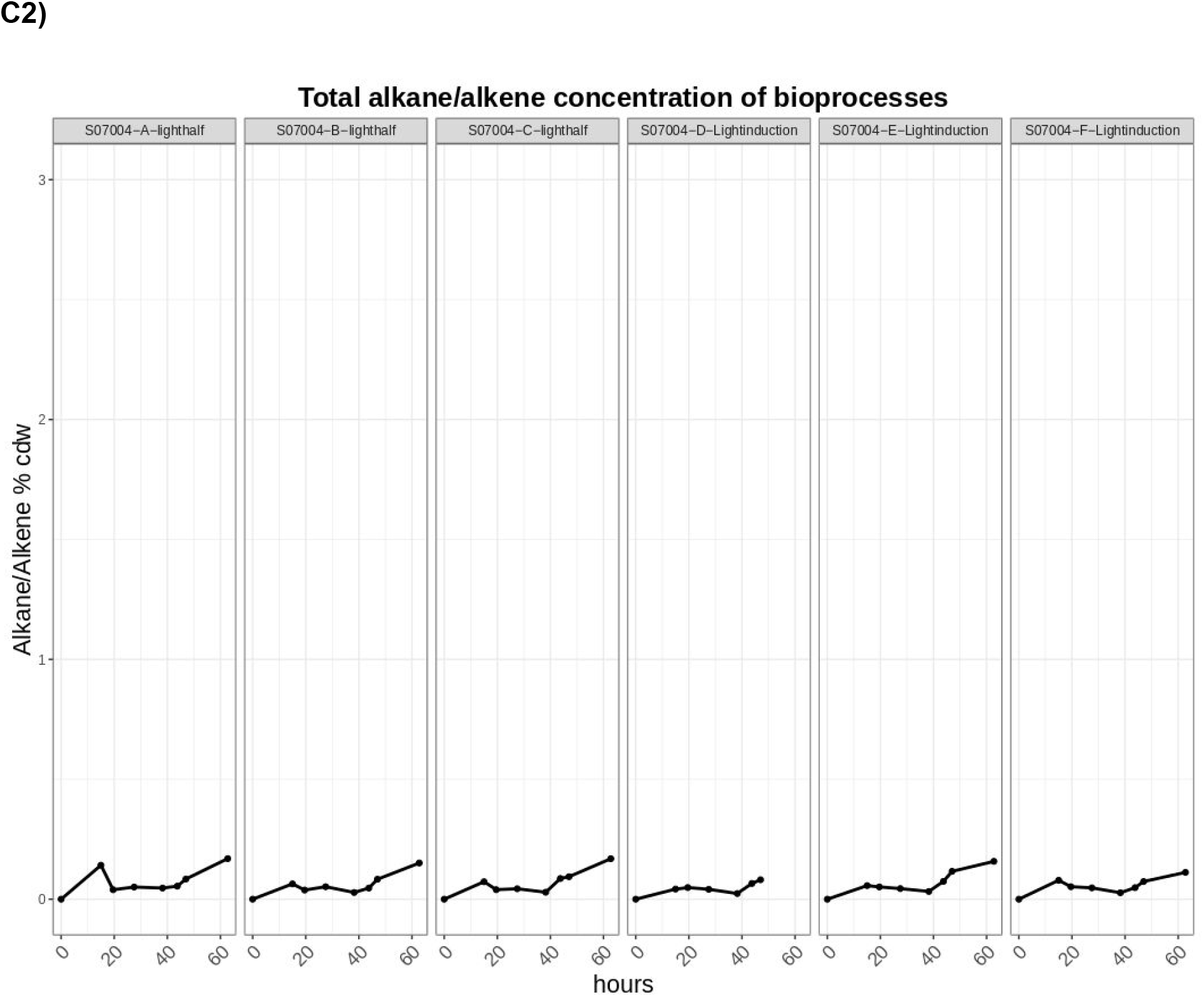
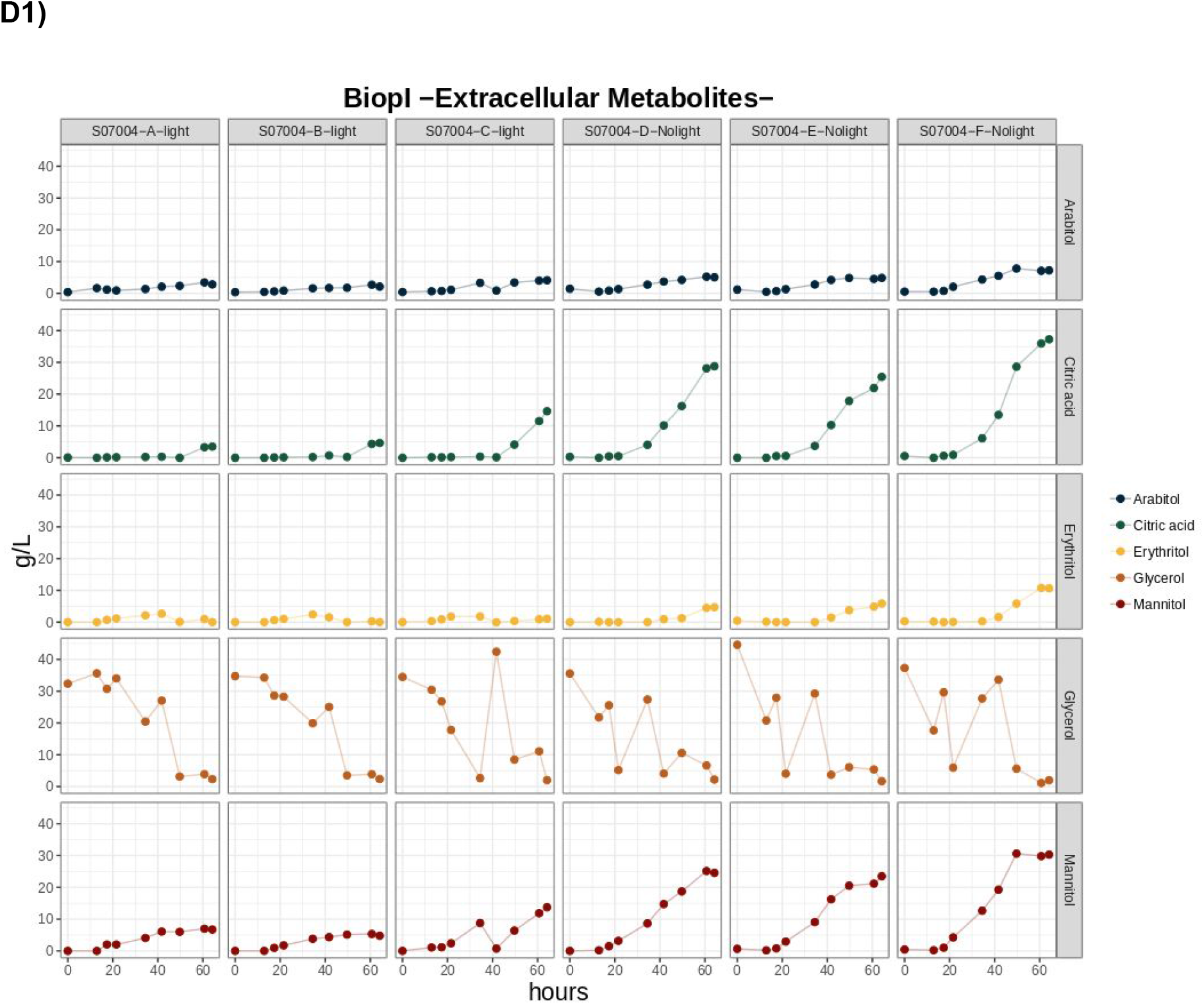
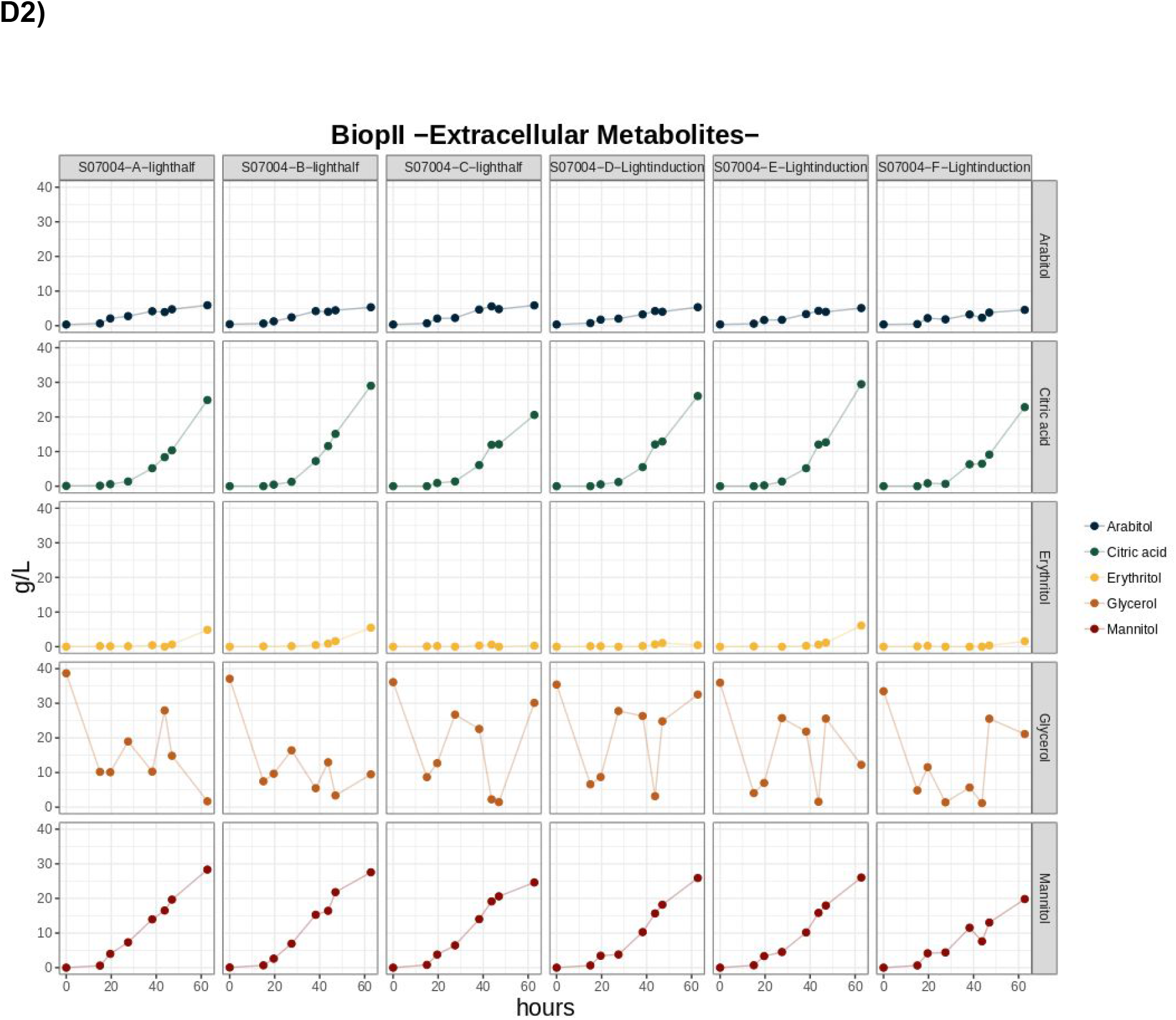

**Fig. S9:**
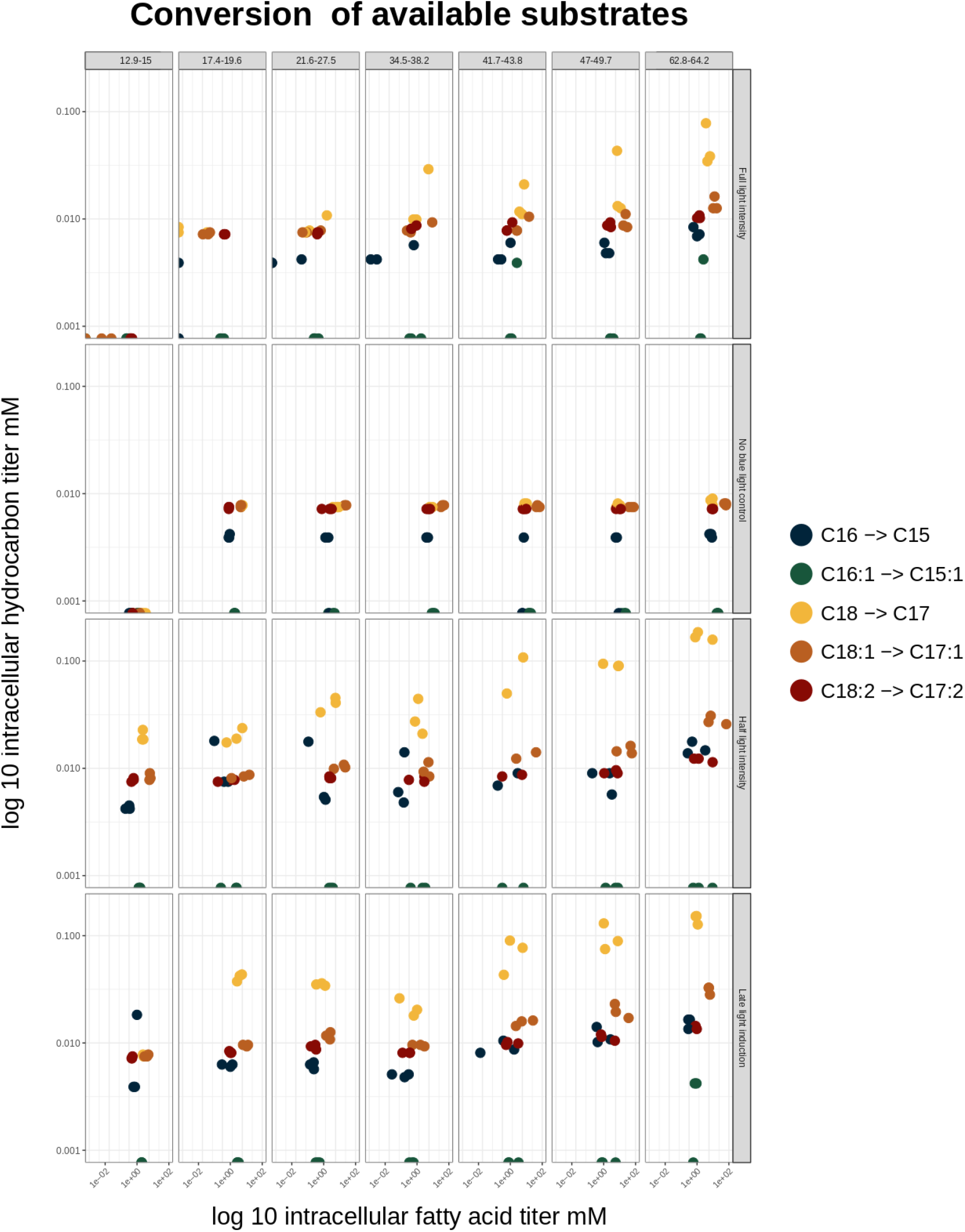

### **Seq 1: Sequence of truncated CvFAP, optimized for *Y. lipolytica* codon usage** _(ordered by baseclear B.V.)_

_ATGCGAGCCTCTGCTGTCGAGGACATTCGAAAGGTGCTCTCTGACTCCTCTTCCCCCGTCGCCGGACAGAAGTACGACTA CATCCTGGTCGGAGGCGGTACCGCTGCTTGTGTGCTGGCTAACCGACTCTCCGCCGACGGCTCTAAGCGAGTCCTGGTG CTGGAGGCTGGTCCTGACAACACCTCCCGAGACGTGAAGATCCCCGCCGCTATTACCCGACTGTTCCGATCTCCCCTGGA CTGGAACCTCTTCTCTGAGCTGCAGGAGCAGCTCGCTGAGCGACAAATCTACATGGCCCGAGGTCGACTGCTCGGAGGCT CTTCCGCCACCAACGCTACCCTGTACCACCGAGGCGCCGCTGGTGACTACGACGCTTGGGGAGTCGAGGGCTGGTCTTC CGAGGACGTCCTCTCCTGGTTCGTGCAGGCCGAGACCAACGCTGACTTCGGACCTGGTGCTTACCACGGCTCTGGTGGA CCTATGCGAGTCGAGAACCCCCGATACACCAACAAGCAGCTGCACACCGCCTTCTTCAAGGCCGCTGAGGAAGTCGGACT CACCCCCAACTCCGACTTCAACGACTGGTCTCACGACCACGCTGGTTACGGAACCTTCCAGGTCATGCAGGACAAGGGCA CCCGAGCCGACATGTACCGACAGTACCTGAAGCCCGTCCTCGGTCGACGAAACCTGCAGGTGCTCACCGGAGCCGCTGT CACCAAGGTGAACATTGACCAGGCTGCTGGAAAGGCTCAGGCTCTGGGCGTCGAGTTCTCCACCGACGGTCCCACCGGA GAGCGACTGTCCGCTGAGCTCGCTCCCGGCGGTGAGGTCATTATGTGTGCTGGTGCTGTGCACACCCCCTTCCTGCTCAA GCACTCTGGTGTGGGTCCTTCTGCTGAGCTGAAGGAGTTCGGTATCCCCGTCGTGTCCAACCTCGCTGGAGTGGGACAGA ACCTGCAGGACCAGCCTGCTTGTCTCACCGCTGCTCCCGTGAAGGAGAAGTACGACGGAATCGCCATTTCTGACCACATC TACAACGAGAAGGGCCAGATTCGAAAGCGAGCCATCGCTTCCTACCTGCTCGGAGGCCGAGGTGGACTGACCTCTACCG GCTGTGACCGAGGTGCCTTCGTCCGAACCGCTGGTCAGGCTCTGCCCGACCTCCAGGTCCGATTCGTGCCTGGAATGGC TCTCGACCCCGACGGTGTCTCCACCTACGTGCGATTCGCCAAGTTCCAGTCCCAGGGTCTGAAGTGGCCCTCTGGAATTA CCATGCAGCTCATCGCCTGTCGACCCCAGTCCACCGGCTCTGTGGGTCTGAAGTCTGCCGACCCCTTCGCCCCTCCCAAG CTCTCTCCTGGATACCTGACCGACAAGGACGGTGCCGACCTGGCTACCCTCCGAAAGGGTATTCACTGGGCTCGAGACGT GGCTCGATCTTCTGCTCTGTCCGAGTACCTCGACGGAGAGCTGTTCCCCGGTTCCGGAGTCGTGTCTGACGACCAGATCG ACGAGTACATTCGACGATCTATCCACTCTTCCAACGCTATCACCGGTACCTGTAAGATGGGAAACGCCGGCGACTCTTCCT CTGTCGTGGACAACCAGCTGCGAGTCCACGGTGTGGAGGGACTCCGAGTCGTGGACGCCTCTGTCGTTCCTAAGATTCCC GGCGGTCAGACCGGAGCTCCTGTCGTGATGATTGCTGAGCGAGCCGCTGCCCTGCTCACTGGCAAGGCTACTATCGGTG CTTCCGCCGCCGCTCCTGCTACTGTGGCTGCTTAG_

### Anova1

Coefficients:

Estimate Std. Error t value Pr(>|t|)

(Intercept) −5.426e-19 2.935e-04 0.000 1.0000

ConditionFull-Intensity & Pulse 0ms 9.805e-03 5.359e-04 18.296 3.76e-12 ***

ConditionFull-Intensity & Pulse 100ms 6.939e-03 5.359e-04 12.949 6.78e-10 ***

ConditionFull-Intensity & Pulse 5000ms 1.881e-03 5.359e-04 3.511 0.0029 **

ConditionHalf-Intensity & Pulse 0ms 7.898e-03 5.359e-04 14.738 9.95e-11 ***

ConditionHalf-Intensity & Pulse 100ms 7.100e-03 6.227e-04 11.403 4.29e-09 ***

ConditionHalf-Intensity & Pulse 5000ms 9.154e-04 6.227e-04 1.470 0.1609

### Anova2

Coefficients:

Estimate Std. Error t value Pr(>|t|)

(Intercept) 0.0001653 0.0004361 0.379 0.7100

ConditionFull-Intensity & Pulse 0ms 0.0087410 0.0007553 11.573 7.08e-09 ***

ConditionFull-Intensity & Pulse 100ms 0.0080151 0.0007553 10.612 2.27e-08 ***

ConditionFull-Intensity & Pulse 5000ms 0.0018260 0.0007553 2.418 0.0288 *

ConditionHalf-Intensity & Pulse 0ms 0.0085443 0.0007553 11.313 9.63e-09 ***

ConditionHalf-Intensity & Pulse 100ms 0.0053621 0.0008721 6.149 1.86e-05 ***

ConditionHalf-Intensity & Pulse 5000ms 0.0009589 0.0008721 1.100 0.2889

### Sequence for DO-dependent automated feeding using software IRIS v6.0 (6.0.1054.817) from Infors HT, Switzerland

#0, DO dropping,10

IF(pO2.v<40){SEQ=1}

#1,Check feeding volume,1

IF(Feed_Pump.v>100){Feed.sp=0 AND SEQ=0}ELSE{SEQ=2}

#2,DO Check,10

IF(pO2.v>70){Feed.sp=100 AND SEQ=3}ELSE{Feed.sp=0}

#3,Feeding,1

IF(SEQ_TIME>XXX dependent of pump performance XXX){Feed.sp=0;SEQ=4}

#4,Pause,10 IF(SEQ_TIME>Time(1:0)){SEQ=1}

The PyMOL Molecular Graphics System, Version 2.0 Schrödinger, LLC.

